# Antibody Blockade of Ly49/MHC-I interactions enhances Innate and Adaptive Immunity Against Cancer Metastasis

**DOI:** 10.64898/2026.05.07.722994

**Authors:** Abir K. Panda, Surajit Sinha, Kannan Natarajan, Jiansheng Jiang, Sruthi Chempati, Soha Kazmi, Yong-Hee Kim, Suveena Sharma, Paul Schaughency, Lisa F. Boyd, Jonathan M. Hernandez, David H. Margulies, Ethan M. Shevach

## Abstract

**Background:** Antibody-mediated blockade of innate receptor–MHC-I interactions represents a promising strategy to enhance anti-tumor immunity, particularly against metastatic cancers resistant to conventional checkpoint inhibitors. In this study, we investigated the effects of the pan anti-MHC-I monoclonal antibody M1/42, which targets MHC-I interactions with Ly49, selectively expressed on murine NK cell subsets.

**Methods:** We administered M1/42 to mice and assayed the proliferation and activation immune cells. Anti-tumor activity of growth and metastasis of checkpoint inhibitor-resistant pancreatic ductal adenocarcoma (PDAC) and B16F10 melanoma were assessed, complemented by extensive cellular phenotypic and RNA expression analysis. Binding and cryo-electron microscopic (cryo-EM) and X-ray crystallographic structural studies of M1/42 complexed with the mouse MHC-I molecule, H2-D^d^, examined the Ab interaction site in comparison with those of Ly49 inhibitory receptors.

**Results:** M1/42 administration in mice robustly unleashed the proliferation and activation of natural killer (NK) cells, memory CD4^+^ and CD8^+^ T cells, dendritic cells, and macrophages in both lymphoid and non-lymphoid tissues, independent of Fcγ receptors. M1/42 significantly restricted the growth and metastasis of checkpoint inhibitor–resistant pancreatic ductal adenocarcinoma (PDAC) and B16F10 melanoma in the liver and lungs, accompanied by increased tumor infiltration of effector CD8^+^ T cells, reduction of T regulatory cells, and a pro-inflammatory cytokine milieu. The anti-tumor effects of M1/42 depend on NK cells and are associated with upregulation of genes involved in antigen processing, interferon gamma responsiveness, and Th1 cytokine production, while downregulating inhibitory PD1/11 signaling. Structural analysis indicated that the effect of M1/42 on Ly49/MHC-I interactions was not due to direct steric competition.

**Conclusions:** Collectively, these findings demonstrate that M1/42 unleashes coordinated innate and adaptive immune responses, overcoming tumor-induced immunosuppression and resistance to checkpoint blockade. This approach represents a paradigm shift in cancer immunotherapy, offering potential for more effective treatment of metastatic cancers that evade immune surveillance through MHC-I modulation.

**KEY MESSAGES:** *What is already known on this topic:* A pan anti-mouse MHC-I mAb (M1/42) blocks interaction with several NK inhibitory receptors (Ly49A or Ly49C) resulting in NK cell activation and anti-viral and anti-tumor responses *in vitro* and *in vivo*. Other pan anti-human MHC-I mAbs (DX17 and W6/32) function similarly, blocking LILRB inhibitory receptor interaction of myeloid cells and NK cells. These stimulate human immune cells in humanized mouse models.

*What this study adds:* This study analyzes the effects of the pan anti-mouse MHC-I mAb on NK and myeloid cell activation in detail, in the absence of T or B cells, and independent of FcR interaction. Additionally we analyze several mouse models of metastatic tumor progression, indicative of the progressive activation not only of the innate immune response, but also adaptive responses. The molecular mechanism of the mAb blocking of inhibitory receptors is revealed by cryo-EM and X-ray structures of M1/42 Fab/MHC-I (H2-D^d^) complexes.

*How this study might affect research, practice, or policy:* Elucidation of the details of the inhibitory effects of the mouse pan anti-mouse MHC-I mAb provides not only a more advanced understanding of the murine model system, but suggests additional functional avenues to be explored using the parallel an anti-human MHC-I mAbs.

## INTRODUCTION

Major histocompatibility complex class I (MHC-I) molecules are ubiquitously expressed cell surface glycoproteins present on nearly all nucleated cells ^1, 2^. They serve as essential ligands for both adaptive and innate immune cells, mediating the presentation of endogenous peptides to cytotoxic (CD8^+^) T lymphocytes and acting as critical regulators of innate immune responses ^3–7^. Downregulation of MHC-I expression, which may occur during viral infection or neoplastic transformation, leads to the activation of natural killer (NK) cells through the “missing self” mechanism, thereby promoting the elimination of abnormal cells ^7–9^.

Innate immune cells, particularly NK cells, utilize a diverse repertoire of germline-encoded receptors that recognize MHC-I molecules ^10^. In mice, the Ly49 family of C-type lectin-like receptors is predominantly responsible for MHC-I recognition ^11–13^. In humans, functional analogs include the killer cell immunoglobulin-like receptors (KIR), leukocyte immunoglobulin-like receptors (LILR), and the CD94/NKG2A heterodimer ^14^. Both activating and inhibitory isoforms of these receptors are co-expressed on overlapping NK cell subsets, resulting in a finely tuned regulatory network that governs NK cell effector functions ^5, 15, 16^. Comparative genomic and crystallographic studies reveal that, while Ly49 and KIR families evolved independently and are structurally unrelated, both may sense MHC-I polymorphism, demonstrating convergent evolution driven by the need for host immune surveillance ^17–20^. The structural basis for inhibitory signaling is also conserved, as these receptors often recruit phosphatases such as SHP-1 or SHP-2 upon ligand engagement, dampening NK cell activation ^12, 21, 22^.

We have previously reported that the pan anti-mouse MHC-I antibody M1/42 effectively blocks Ly49/MHC-I interactions, unleashing robust NK cell activation and subsequent IFNγ production. This cytokine-driven cascade promotes the expansion of memory phenotype (MP) CD4^+^ and CD8^+^ T cells and enhances the immune response to tumors and viral infections ^23^. We have recently extended this strategy to humans and demonstrated that pan anti-human MHC-I antibodies such as W6/32 and DX17 block LILR/MHC-I interactions, resulting in global NK and myeloid activation and improved tumor control in humanized mouse models of pancreatic cancer ^24^. Additionally, we have shown that another mAb, B1.23.2, that recognizes almost all HLA-A, -B, and -C molecules (except-–HLA-A*02 and -A*68 alleles), induces NK and myeloid cell activation by blocking KIR/MHC-I interactions ^25^. In this report, we demonstrate that blocking Ly49/MHC-I interactions results in activation of NK and myeloid cells in the absence of T and B cells, and that activation does not require Fc receptor (FcR)-mediated crosslinking of the mAb. We then use several well-characterized models of tumor metastatic disease in the mouse and show that pan anti-MHC-I treatment has profound therapeutic effects initiated by activating the innate immune response with subsequent enhancement of the adaptive immune response. Cryo-electron microscopic (cryo-EM) and X-ray structural studies of M1/42/H2-D^d^ complexes offer mechanistic insight to these functions. Taken together, these studies demonstrate that activated NK cells can control tumor metastases in mice and strongly suggest that blocking LILR/HLA interactions in the human ^24^ may have similar therapeutic applications.

## MATERIALS AND METHODS

Detailed description of materials and methods can be found in the supplemental materials and methods (online supplemental file 1).

## RESULTS

### Activation of NK and myeloid cells by blocking Ly49/MHC-I interactions is T-cell and FcR-independent

We have previously demonstrated that administration of M1/42 mAb *in vivo* to C57BL/6 mice for 8 days markedly enhanced splenic NK cell proliferation, accompanied by increased numbers of CD4^+^ effector memory (EM, CD44^+^CD62L^−^), CD8^+^ central memory (CM, CD44^+^CD62L^+^), and CD8^+^ EM activation. To determine whether NK and myeloid cell activation required the presence of T cells, M1/42 was administered to RAG^−/−^ mice. Significantly enhanced NK cell proliferation (indicated by increased Ki-67 staining) and IFNγ production (figure 1A), as well as the proliferation of CD11b^+^ monocytes and CD11c^+^ dendritic cells (figure 1B), were observed *in vivo*. M1/42-mediated proliferation of CD11b^+^ and CD11c^+^ myeloid cells depended on the presence of NK cells and NK-derived IFNγ, as NK cell depletion (with anti-NK1.1) and neutralization of IFNγ completely reversed the enhanced proliferation of myeloid cells induced by M1/42 (figure 1C-F) ^23^. To demonstrate that the effects of M1/42 administration *in vivo* were solely the result of blocking Ly49/H-2 interactions and not mediated by mAb-mediated signaling, FcγR^−/−^ mice were treated with M1/42. Proliferation of NK cells and memory T cells was similar to that seen in wild-type mice, confirming that M1/42-mediated lymphocyte proliferation is FcγR-independent (online supplemental figure S1A-E) ^26^.

**Figure 1.**
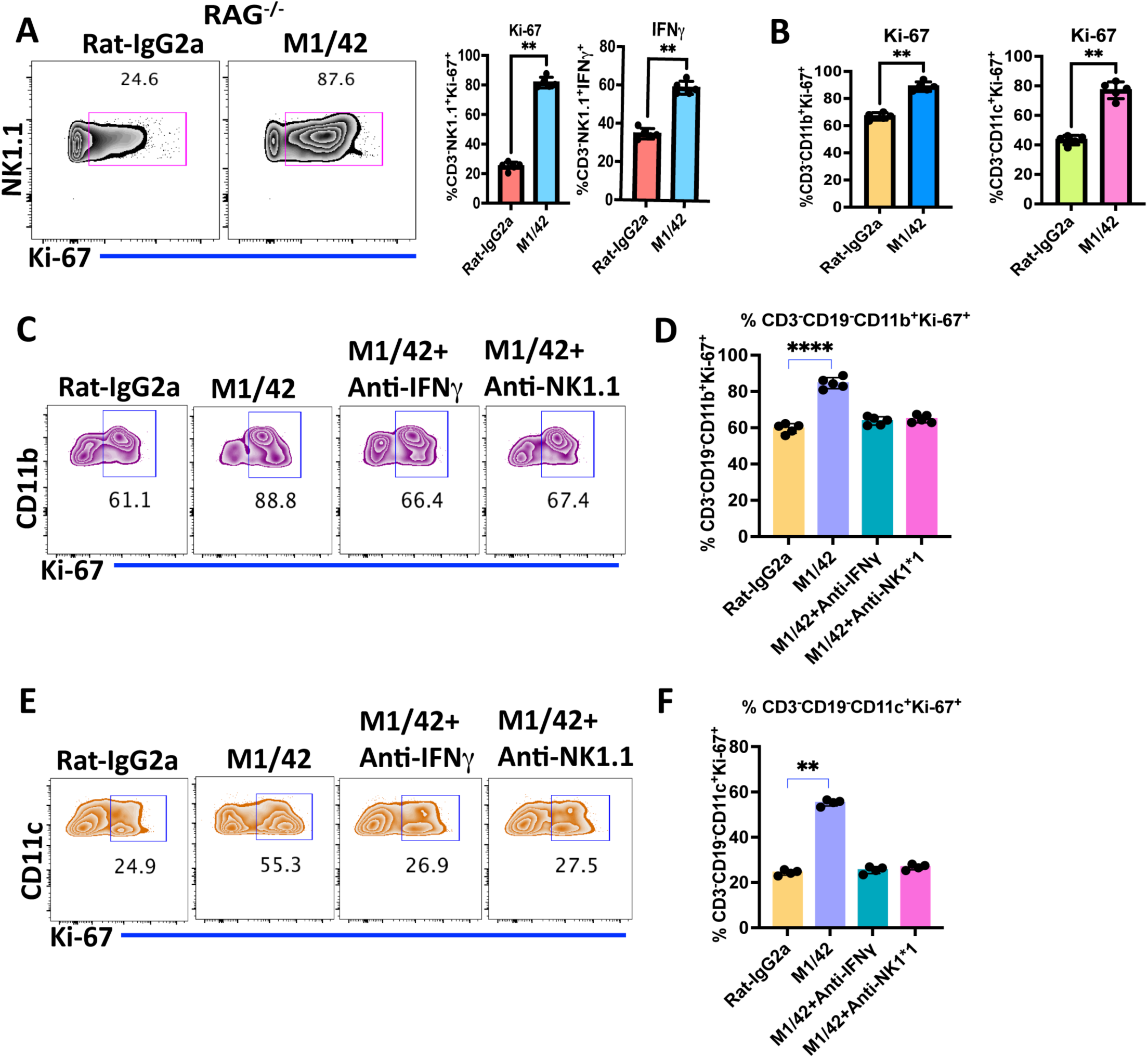
M1/42-mediated NK and myeloid cell proliferation is T cell independent. (A) Flowcytometric contour plots and statistical analysis of NK cell proliferation (Ki-67) and percentage of IFNγ expression of splenocytes from Rag^−/−^ mice, treated with Rat-IgG2a or M1/42 on days 0, 2, 4, and 6, and spleens were harvested on day 8. (B) Percentage of Ki-67^+^ CD3^−^CD11b^+^ monocytes and CD3^−^CD11c^+^ dendritic cells from Rag^−/−^ splenocytes after treatment with rat-IgG2a or M1/42 on day 8. (C-F) C57BL/6 mice were injected with neutralizing anti-IFNγ or NK depleting (anti-NK 1.1) mAb starting from day −2 and day −1 and subsequently cotreated with M1/42 on days 0, 2, 4, and 6, and splenocytes were analyzed on day 8. Flowcytometric Ki-67 analysis of CD3^−^CD11b^+^ monocytes and CD3^−^CD11c^+^ dendritic cells from C57BL/6 derived splenocytes on day 8 after mAb treatments.

### M1/42 administration enhances the proliferation of liver-derived innate and adaptive immune cell subsets

As tumor metastases frequently involve the liver, it was of interest to determine whether the activating properties of M1/42 would also be observed in non-lymphoid sites. M1/42 was administered every other day for 6 days, and liver lymphocytes were analyzed on day 8. M1/42 treatment markedly enhanced NK cell proliferation (figure 2A) however the total frequency of NK cells remained unchanged (figure 2B). M1/42 treatments also enhanced the proliferation of liver resident CD11c^+^XCR1^+^ dendritic cells (figure 2C) and CD11b^+^F4/80^+^ macrophages (figure 2D). M1/42 treatment did not affect the frequency of CD4^+^CD1d^+^ NKT cells (figure 2E) but significantly enhanced CD4^+^CD1d^+^ NKT cell proliferation (from ∼10% to 75%) (figure 2F). M1/42 also modestly enhanced liver CD4^+^CD44^+^CD62L^−^ EM T cell frequency (figure 2G) and proliferation (figure 2H). M1/42 treatments also markedly enhanced CD8^+^CD44^+^CD62L^−^ EM T-cell frequency (15% to 55%) but had no effect on the percentage of CD8^+^CD44^+^CD62L^+^ CM T cells (figure 2I). M1/42 significantly enhanced the proliferation of CD8^+^CM, and CD8^+^ EM T cells (figure 2J-L). Taken together, these studies strongly suggest that the effects of M1/42 result in profound activation of innate and adaptive immune systems at both lymphoid and non-lymphoid sites.

**Figure 2.**
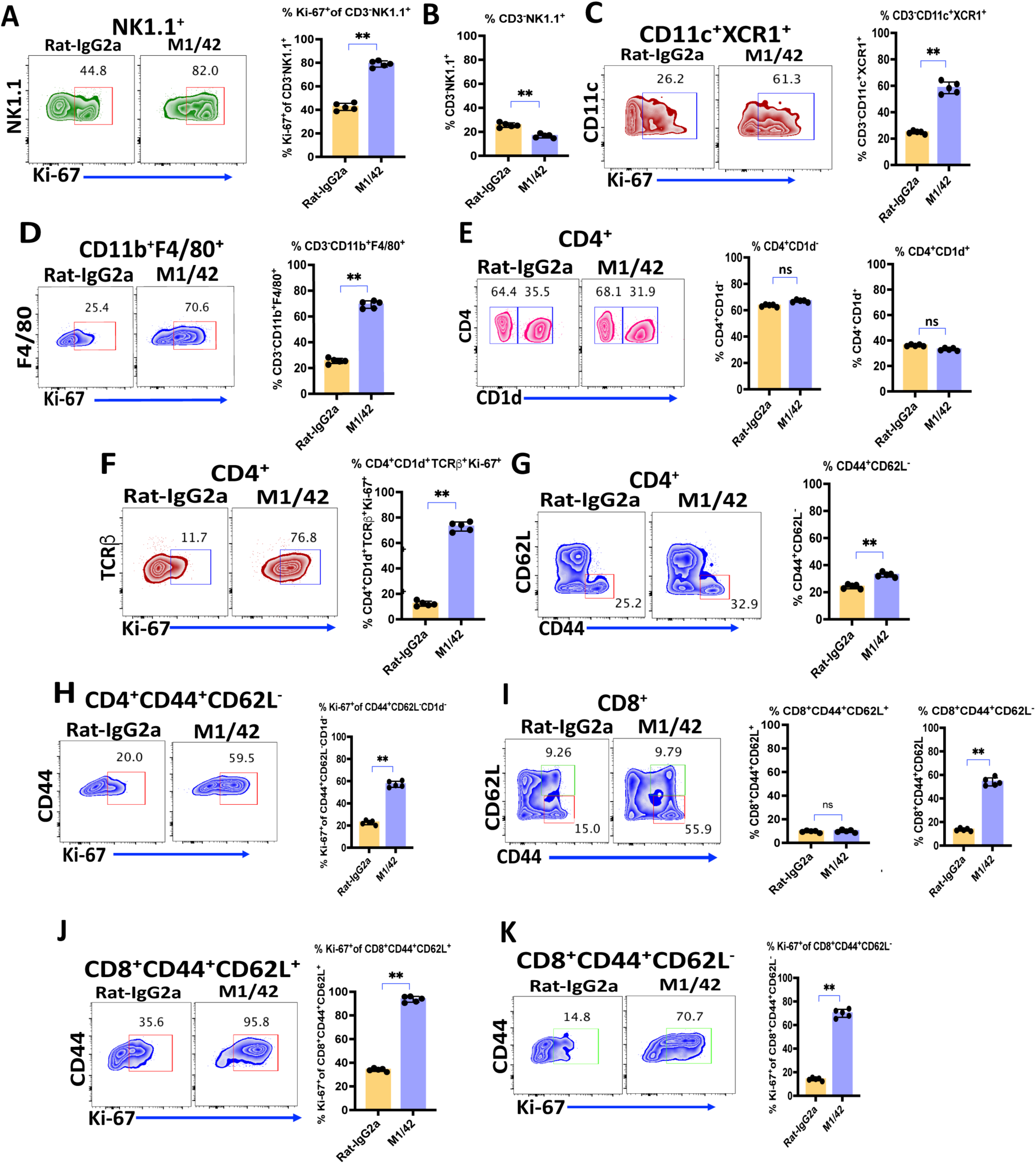
M1/42 enhances homeostatic NK, myeloid, and T cell proliferation in liver lymphocytes. C57BL/6 mice were injected with rat-IgG2a or M1/42 on days 0, 2, 4, and 6, and liver lymphocytes were harvested on day 8. (A & B) Flowcytometric contour plot analysis of proliferation and frequency of NK1.1^+^ NK cell subsets on day 8 after antibody treatment. (C & D) Proliferation of CD11c^+^XCR1^+^ dendritic cells and CD11b^+^F4/80^+^ macrophages on day 8. (E & F) Percentage and proliferation of CD1d^+^CD4^+^ NKT cells on Day 8. (G) Percentage of CD4^+^CD62L^−^CD44^+^ effector memory T cells on Day 8. (H) Percentage of CD8 central memory (CD8^+^CD44^+^CD62L^+^) and CD8 effector memory (CD8^+^CD44^+^CD62L^−^) cells after antibody treatment on day 8. (I, J, & K) Ki-67 expression in CD4^+^CD62L^−^CD44^+^ effector memory, CD8 central memory (CD8^+^CD44^+^CD62L^+^) and CD8 effector memory (CD8^+^CD44^+^CD62L^−^) T cells on day 8.

### M1/42 inhibits the growth of checkpoint inhibitor-resistant KPC cells in pancreatic ductal adenocarcinoma (PDAC) models of tumor metastasis

Our previous studies demonstrated that M1/42 administration significantly restricted the growth of anti-CTLA-4 and anti-PD1 checkpoint inhibitor-sensitive MC38 colon adenocarcinoma. In this model, M1/42-mediated anti-tumor immunity involved both NK cell-mediated innate and CD8^+^ T cell-mediated adaptive immune responses. Depletion of either NK cells or CD8^+^ T cells completely reversed the anti-tumor effects of M1/42. Although both anti-CTLA4 and anti-PD1 limit MC38 colon adenocarcinoma in mice, these therapies are effective in only 10-20% of primary or metastatic cancer patients with survival beyond five years. Checkpoint inhibitor therapies have also not produced reliable clinical benefits in human PDAC ^27^. We evaluated the anti-tumor effects of M1/42 in the KPC (*Kras^G12D/+^, Trp53^R^*^172*H/+*^*, Pdx-1-Cre*) subcutaneous mouse model of PDAC. Activating point mutations in the KRAS protooncogene (G12D, present in 80%-90% of cases) and dominant negative mutations in the Trp53 tumor suppressor gene (R172H, present in 50%-75% of cases) are the most common genetic alterations in human PDAC. KPC tumor cells were injected subcutaneously into C57BL/6 mice, followed 3 days later by M1/42 mAb treatment twice weekly for three weeks, starting on day 3. This regimen significantly restricted tumor development, with 40% of animals exhibiting no tumor growth (online supplemental figure S2A, B).

The anti-tumor effects of M1/42 were then investigated in PDAC metastatic models. Fluorescent KPC cells were injected into the spleen capsule, followed by splenectomy 5 minutes later, allowing tumor cell migration to the liver (figure 3A). M1/42 was administered twice weekly from day 3 to day 20, and metastatic outgrowth and liver colonization were analyzed on days 0, 14, and 21 using IVIS bioluminescence *in vivo* whole-body imaging (online supplemental figure S2C, D). Metastatic outgrowth of KPC cancer cells on day 22 was significantly lower in the livers of the M1/42-treated group compared to the isotype control group (online supplemental figure S2B). The fluorescence photon efflux remained similar on day 14 but was markedly reduced on day 21 following M1/42 treatment. Bioluminescence imaging and statistical analysis of quantitative photon efflux of harvested liver on day 22 further confirmed that M1/42 treatment markedly restricted KPC cancer cell-derived liver metastasis in the PDAC model (online supplemental figure S2E & S2F). Flow-cytometric analysis of tumor-infiltrated lymphocytes (TILs) from metastatic liver revealed marked enhancement (∼ 2 to 4-fold) of the frequency of CD4^+^ and CD8^+^ T cells in the M1/42 treatment group (online supplemental figure S2G), decreased percentages of CD4^+^Foxp3^+^ Treg (online supplemental figure S2H), but significantly enhanced frequency of CD4^+^ and CD8^+^ memory T cells in the TILs (online supplemental figure S2I, J). CD11c^+^ dendritic cells also increased 2-3-fold while the frequency of B cells (CD19^+^) was significantly lower in the M1/42-treated group in TILs from metastatic liver (online supplemental figure S2K). Mature and functional NK cells (*Tbet^+^Eomes^+^*, ^28, 29^) were significantly increased in the livers following M1/42 treatment. Furthermore, M1/42 treatment enhanced the co-expression of IFNγ and TNFα and increased the levels of Granzyme B and CXCR3 expression on NK1.1^+^ NK cells (figure 3C, D). M1/42 treatment also upregulated co-expression of the transcription factors Tbet and Eomes, the expression of IFNγ and TNFα, as well as Granzyme B and CXCR3 in MP CD4^+^ and CD8^+^ T cells (figure 3E-H).

**Figure 3.**
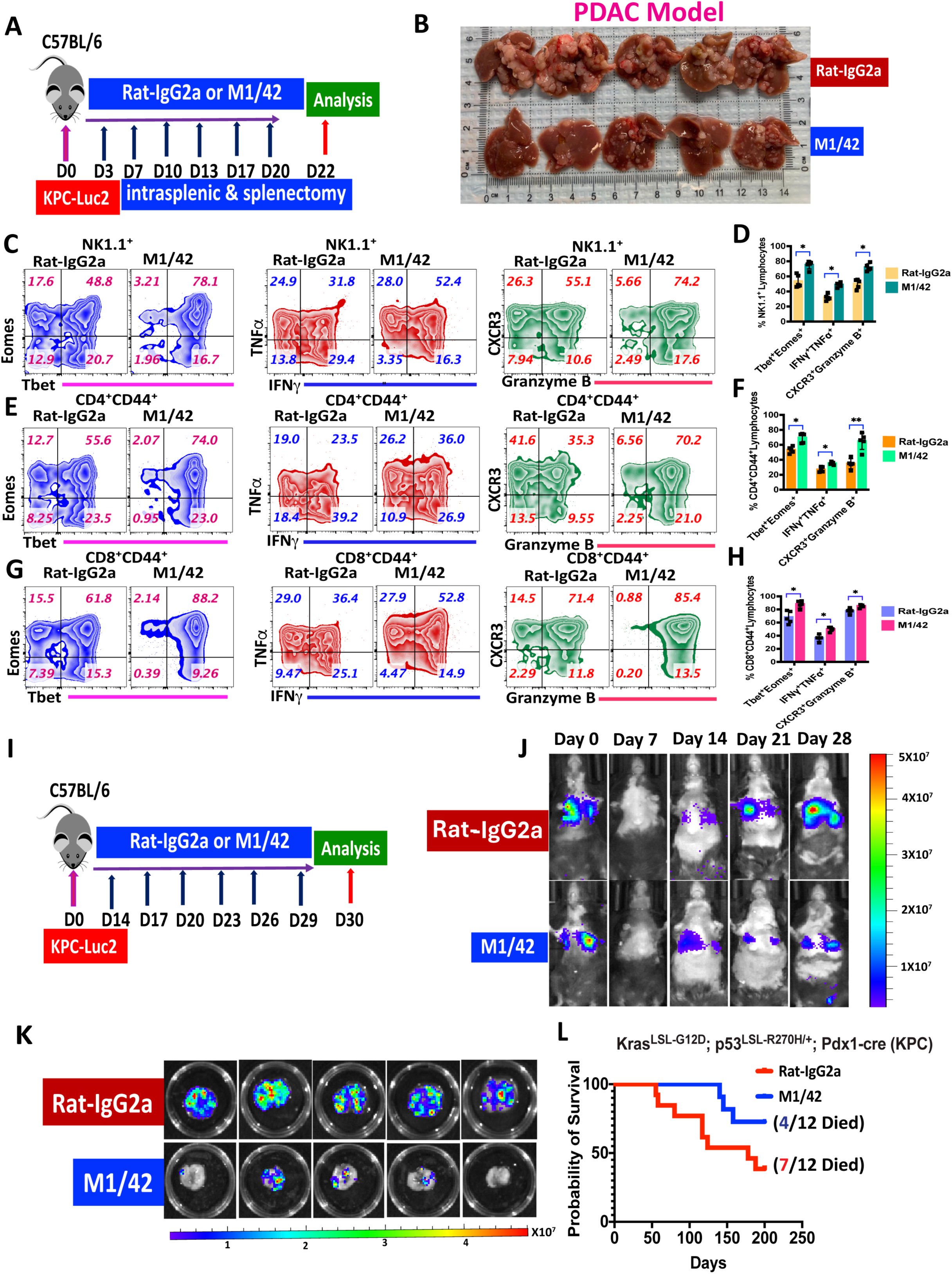
M1/42 administration unleashes anti-tumor immunity to checkpoint inhibitor-resistant KPC cancer cells harboring *KRAS^G12D^ and TP53^R172H^* point mutations in pancreatic ductal adenocarcinoma (PDAC) metastasis model. (A) KPC-Luc-2 cell lines were injected intrasplenically in C57BL/6 mice, followed by splenectomy to establish liver metastasis, rat-IgG2a or M1/42 antibodies were injected i.p. at 3 day intervals from day 3 to day 20, and the livers were harvested on day 22. (B) KPC cell outgrowth in the liver after antibody treatments on day 22. (C & D) Flowcytometric contour plot analysis of liver-derived NK cells for percentage of Tbet^+^Eomes^+^, TNFα^+^IFNγ^+^, and CXCR3^+^Granzyme B^+^ expression. (E & F) Flowcytometric contour plot analysis of metastatic liver-derived CD4^+^CD44^+^ memory T cell for percentage for Tbet^+^Eomes^+^, TNFα^+^IFNγ^+^, and CXCR3^+^Granzyme+ expression. (G & H) Flowcytometric contur plot analysis of metastatic liver-derived CD8^+^CD44^+^ memory T cells for percentage for Tbet^+^Eomes^+^, TNFα^+^IFNγ^+^, and CXCR3^+^Granzyme-B^+^ expression. (I) KPC-Luc-2 cells were injected into the tail vein of C57BL/6 animals to establish lung metastasis, rat-IgG2a or M1/42 antibody was injected i.p. at 3 day intervals from day 14 to day 28 and lungs were harvested on day 30. (J) Representative images of weekly *in vivo* bioluminescence to determine metastatic outgrowth after rat-IgG2a or M1/42 treatment from day 14. (K) Pulmonary outgrowth of KPC cells was detected by bioluminescence on day 30 after rat-IgG2a and M1/42 treatment (5 mice per group) at 3 day intervals starting from day 14. (L) Survival growth curve of spontaneously developed tumors in KPC (*LSL-KrasG12D/+; LSL-Trp53R172H/+; Pdx-1-Cre)* mice after Rat-IgG2a and M1/42 treatment once a week, from week 7 to week 16.

The ability of M1/42 treatment to inhibit the growth of PDAC metastases in the liver also extended to the lung. Fluorescent KPC cells were injected into the tail veins of C57BL/6 mice, which were then monitored for lung metastatic outgrowth by IVIS whole body imaging at weekly intervals (figure 3I-J). The bioluminescent KPC cells appeared to be completely eliminated on day 7, likely secondary to clearing by the innate immune system, but a population resistant to immune attack became visible at day 14. M1/42 treatment from day 14 to day 28 significantly restricted the growth of the newly emergent KPC cells in the lung, as shown by bioluminescence of whole body imaging and harvested lungs (figure 3J, K). Analysis of TILs from lung metastases revealed that the frequency of CD8^+^ T cells was twice the frequency of CD4^+^, with a modest reduction in Treg (CD4^+^FoxP3^+^) among the TILs from treated mice (online supplemental figure S3A, B). The frequencies and proliferation of CD4^+^ and CD8^+^ EM T cells were markedly enhanced after M1/42 treatment (online supplemental figure S3C-F). M1/42 treatment also increased the percentages and proliferation of NK cells and decreased the frequency of CD11b^+^Gr1^+^ myeloid suppressor cells in the lung tumor microenvironment (online supplemental figure S3G-I). Lastly, M1/42 treatment of mice that expressed Kras and p53 mutations (*LSL-Kras^G12D/+^; LSL-Trp53^R172H/+^; Pdx-1-Cre* mice) significantly enhanced their survival in a spontaneous PDAC tumor model (figure 3L). These results strongly suggest that M1/42 treatment is capable of enhancing local anti-tumor immunity of checkpoint inhibitor-resistant PDAC, liver and lung metastatic outgrowth, and the development of spontaneous tumors in a tumor-susceptible strain.

### M1/42 inhibits checkpoint inhibitor-resistant B16F10 lung metastasis by enhancing NK cytotoxicity and expanding melanoma-specific CD8^+^ T cells

Previous studies have shown that M1/42 treatment significantly inhibited the growth of checkpoint inhibitor-resistant B16F10 melanoma when inoculated subcutaneously ^23^, and others evaluated the effect of M1/42 on lung metastasis of B16F10 ^30^. To further evaluate the effectiveness of M1/42 in a lung metastasis context, B16F10 melanoma cells were injected into the tail vein of mice. M1/42 was administered every three days starting from day 3 until day 18, and tumor-bearing lungs were analyzed on day 20. This treatment regimen resulted in a significant reduction in B16F10 melanoma growth and colonization within the lungs (online supplemental figure S4A, B). Immunohistochemical analysis of lungs infiltrated by B16F10 cells revealed a marked increase in the frequency of CD8^+^ T cells following M1/42 treatment on day 15 (online supplemental figure S4C). To elucidate the roles of NK cells and T cells in controlling B16F10 lung metastasis, mice were depleted of NK and T cells prior to B16F10 melanoma cell engraftment and subsequent M1/42 administration. Depletion of either CD8^+^ T cells alone or both CD4^+^ and CD8^+^ T cells had minimal impact on lung metastasis after two weeks. In contrast, NK cell depletion led to rapid, unchecked progression of melanoma in the lungs and significant systemic morbidity (figure 4 A, B). These results demonstrate that NK cells are essential for M1/42-mediated therapy of B16F10 lung metastases ^31^, although inhibition of tumor growth also requires the adaptive immune response.

**Figure 4.**
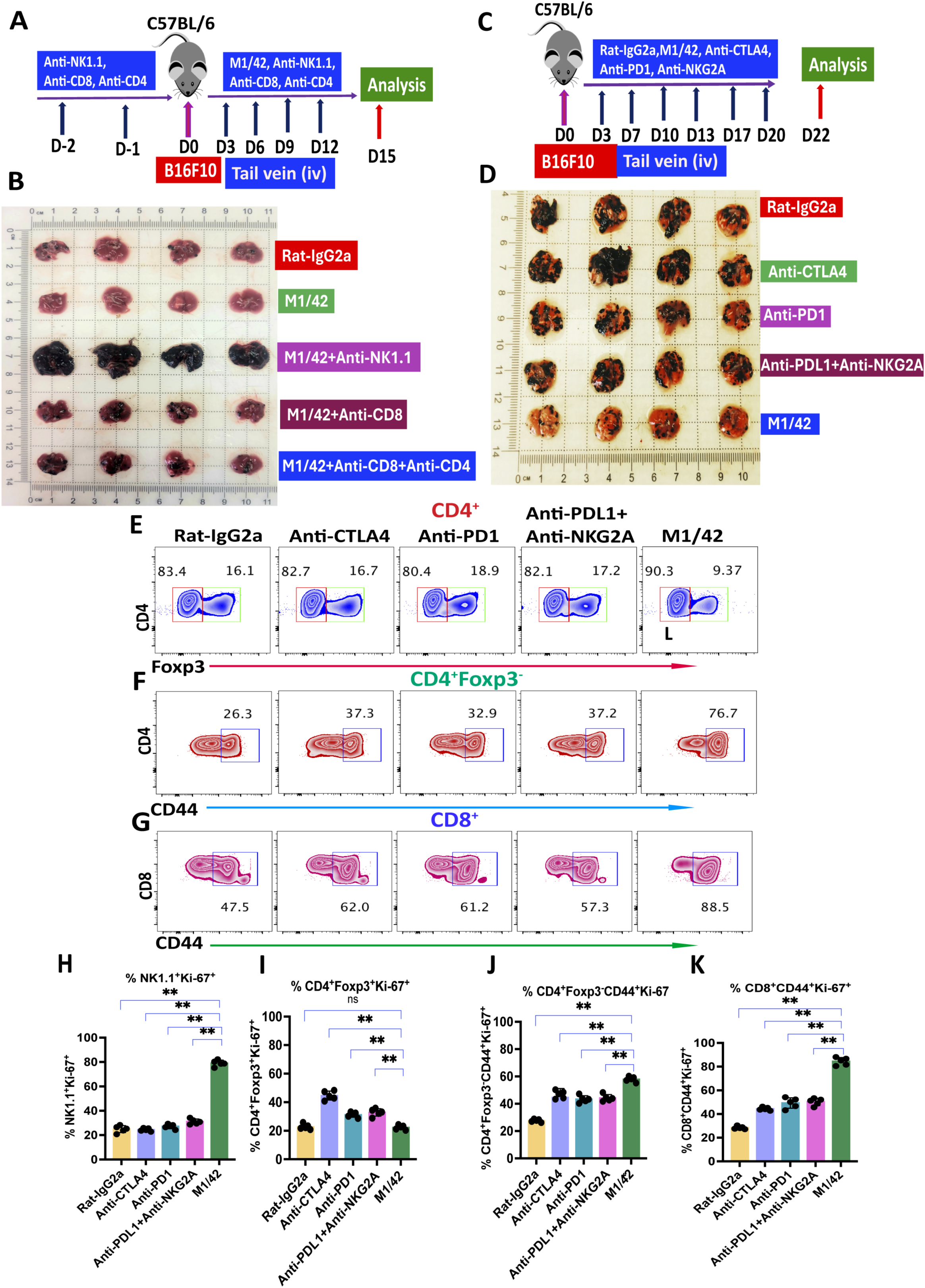
M1/42 administration augments NK-mediated anti-metastatic immunity in checkpoint inhibitor-resistant B16F10 lung metastasis model. (A) C57BL/6 mice received intraperitoneal injections of NK, CD4, and CD8-depleting monoclonal antibodies on days −1 and −2. B16F10 melanoma cells were injected into the tail vein on day 0, followed by administration of M1/42 alone or in combination with anti-NK1.1, anti-CD8, or anti-CD4 and anti-CD8 on days 3, 6, 9, and 12. Lungs were harvested on day 15. (B) B16F10 melanoma cell outgrowth in lungs was assessed on day 15 following antibody treatments. (C) C57BL/6 mice were injected with B16F10 melanoma cells in the tail vein on day 0, followed by administration of rat-IgG2a, M1/42, anti-CTLA4, anti-PD1, or anti-PDL1 with anti-NKG2A every 3–4 days from day 3 to day 20. Lungs were harvested on day 22. (D) Pulmonary metastatic outgrowth of B16F10 melanoma cells was evaluated after treatment with M1/42, anti-CTLA4, anti-PD1, anti-PDL1, and anti-NKG2A, with lungs harvested on day 22. Percentage of (E) CD4^+^Foxp3^+^ Tregs, (F) CD4^+^Foxp3^−^CD44^+^ T cells and (G) CD8^+^CD44^+^ T cells. Ki-67 expression in (H) CD3^−^NK1.1^+^ NK cells, (I) CD4^+^Foxp3^+^ Treg cells, (J) CD4^+^Foxp3^−^CD44^+^ effector T cells and (K) CD8^+^CD44^+^ memory T cells.

To compare directly the efficacy of M1/42 with that of classical checkpoint inhibitors, mice were treated with anti-CTLA4, anti-PD1, anti-PD-L1 plus anti-NKG2A, or M1/42 every 3 to 4 days starting on day 3 for a total of 21 days following B16F10 tumor cell engraftment into the lungs (figure 4C, D). Analysis on day 22 revealed that, compared to all classical checkpoint inhibitors, M1/42 treatment significantly restricted the development of lung metastases (figure 4D). Further evaluation of tumor-infiltrating lymphocytes (TILs) from the lungs demonstrated that M1/42 treatment led to a modest decrease in the frequency of Tregs, while frequencies of both memory CD4^+^ and CD8^+^ T cells were markedly increased relative to other classical checkpoint inhibitor treatments (figure 4E and online supplemental figure S4D-F). In addition, M1/42 significantly enhanced the proliferation of NK cells in the lungs, with proliferation rates rising from 20% to 80% compared to other checkpoint inhibitor regimens (figure 4H). Notably, Treg proliferation was most significantly elevated after anti-CTLA4 administration, and was also modestly increased by anti-PD1, or anti-PD-L1 plus anti-NKG2A treatments (figure 4I). In contrast, M1/42 treatment led to a robust increase in the proliferation of memory T cell subsets derived from metastatic B16F10 lungs, with CD4^+^ memory T cell proliferation increasing threefold (from 20% to 60%) and CD8^+^ memory T cell proliferation increasing fourfold (from 20% to 80%) compared to all other checkpoint inhibitor groups (figure 4J, K).

Suppression of B16F10 tumor outgrowth following M1/42 treatment was closely associated with increased frequencies of melanoma-specific memory CD8^+^ T cells. Specifically, there was a significant elevation in the percentage of H2-D^b^-restricted CD8^+^CD44^+^GP100^+^ and H2-K^b^-restricted CD8^+^CD44^+^Trp2^+^ T cells in comparison to treatment with anti-CTLA4, anti-PD1, and anti-PD-L1 plus anti-NKG2A checkpoint inhibitors (figure 5A, B). M1/42 administration also significantly enhanced the co-expression of IFNγ and Granzyme B in both CD8^+^CD44^+^GP100^+^ and CD8^+^CD44^+^Trp2^+^ melanoma-specific memory CD8^+^ T cells, compared to other checkpoint inhibitor treatments (figure 5C, D & online supplemental figure S4G, H). The total numbers of the melanoma-specific CD8^+^ memory T cells were markedly increased in harvested B16F10 metastatic lungs following M1/42 treatment relative to other checkpoint inhibitors (figure 5E, F). Importantly, the expansion of melanoma-specific memory CD8^+^CD44^+^GP100^+^ and CD8^+^CD44^+^Trp2^+^ T cells were also observed in the splenocytes of tumor-bearing mice after M1/42 treatment (figure 5G, H). Flow-cytometry-based multiplex cytokine assays performed on day 22 revealed that M1/42 treatment, compared to anti-CTLA4, anti-PD1, or anti-PD-L1 with anti-NKG2A, led to a marked increase in Th1-type cytokines, including IL-2, IL-12, IL-18, IFNα, IFNγ, TNFα, and GM-CSF. There was also a modest elevation in IL-6, IL-4, IL-10, and IL-17 levels in the serum of B16F10 lung melanoma-bearing animals (figure 5I). Collectively, these findings demonstrate that M1/42 treatment not only amplifies the innate immune response in the lung but also significantly expands the adaptive, tumor antigen-specific response both locally in the lung and systemically.

**Figure 5.**
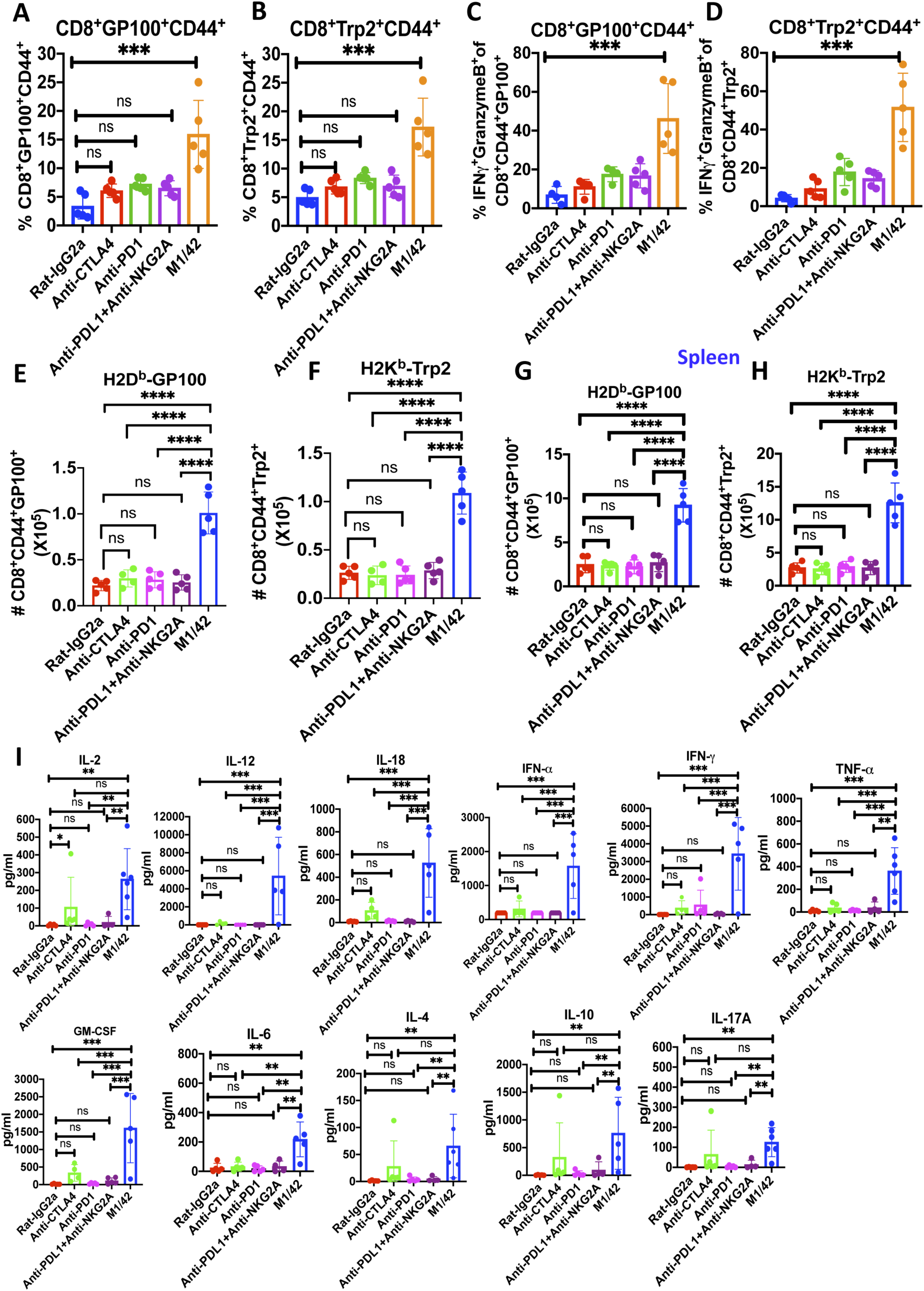
M1/42 administration enhances the expansion of melanoma-specific memory CD8 T cells in the checkpoint inhibitor-resistant B16F10 lung metastasis model. (A & B) Percentage of H2D^b^-GP100^+^ and H2K^b^-TRP2^+^ melanoma-specific CD8^+^CD44^+^ T cells in metastatic lungs were measured on day 22 after treatment with M1/42 and checkpoint inhibitors (anti-CTLA4, anti-PD1, or anti-PDL1 and anti-NKG2A). (C & D) Percentage of IFNγ^+^Granzyme-B^+^GP100^+^ and IFNγ^+^Granzyme-B^+^TRP2^+^ melanoma-specific CD8^+^ T cells in metastatic lungs, determined on day 22 after M1/42 and checkpoint inhibitor treatment. (E & F) Relative total number of H2-D^b^-GP100^+^ and H2-K^b^-TRP2^+^ melanoma-specific CD8^+^ T cells in lungs on Day 22 after M1/42 or other checkpoint inhibitor antibody treatment. (G & H) Relative frequency of H2-D^b^-GP100^+^ and H2-K^b^-TRP2^+^ melanoma-specific CD8^+^ T cells in splenocytes, measured on day 22 after M1/42 and checkpoint inhibitor treatment in the established B16F10 pulmonary metastasis model. (I) Serum cytokines measured by LEGENDplex^TM^ bead array on day 22 in B16F10 lung metastasis mice treated with M1/42 or other checkpoint inhibitors (anti-CTLA4, anti-PD1, or anti-PDL1 with anti-NKG2A) as indicated.

### Gene expression analysis of NK cells from metastatic lungs following M1/42 treatment demonstrates activation of multiple IFN signaling pathways

To investigate the impact of M1/42 treatment on NK cell function, B16F10 was injected as in figure 4C, M1/42 was administered on days 3, 6, and 9 and lung tissue collected on day 10. CD3^−^CD19^+^CD11b^−^CD11c^−^NK1.1^+^ cells sorted from B16F10 lungs and Gene expression profiles were determined using bulk RNA sequencing ^32^. Uniform Manifold Approximation and Projection (UMAP) analysis identified seven distinct, non-overlapping gene clusters in metastatic lungs from B16F10 cells on day 10 after treatment with either Rat-IgG2a or M1/42. Clusters 0, 1, and 2 (figure 6A) were found to be more prevalent in the M1/42 treated mice than clusters 3, 4, 5, and 6 (figure 6B). In a combined analysis of seven clusters on day 10, NK cells from M1/42-treated B16F10 lungs exhibited upregulation of genes such as *Tap1*, *Canx*, *Calr*, *Pdia3, B2m, and Gzmb*. These genes are involved in MHC-I antigen processing and presentation pathway as well as the endoplasmic reticulum (ER) phagosome pathway, the innate immune system, the cellular response to heat stress, and the HSP90-mediated chaperone cycle (figure 6C and online supplemental figure S5A). In contrast, the isotype control group showed upregulation of genes related to signal transduction, mRNA metabolism, and pre-RNA splicing (figure 6D and online supplemental figure S5B). Differential gene expression analysis on all seven clusters by KEGG, GO, and Reactome pathway analyses of clusters 0, 1, and 2 indicated that M1/42 treatment upregulates genes associated with the immune system, antigen processing and presentation (*Tap1, Canx, Calr*), interferon gamma responsiveness, and Th1 cytokines (*Gzma, Gzmb, Prf1, and Nkg7*). Conversely, the control group demonstrated upregulation of genes involved in signal transduction, mRNA metabolism, processing, and splicing (online supplemental figure S5C,D, online supplemental figure S6, and online supplemental figure S7). Furthermore, Ingenuity pathway analysis of differential gene expression showed that M1/42 treatment enhances IFNγ and IFNα/β signaling, IL-15 production, PI3K/Akt signaling, Th1 signaling, IL-12 signaling, and FLT3 signaling, while downregulating the PD1/PD-L1 inhibitory signaling pathway (figure 6E). Taken together, the gene expression analysis of NK cells from M1/42 treated mice demonstrates marked elevation of multiple pathways of NK cells activation, cytokine production and cytokine responsiveness.

**Figure 6.**
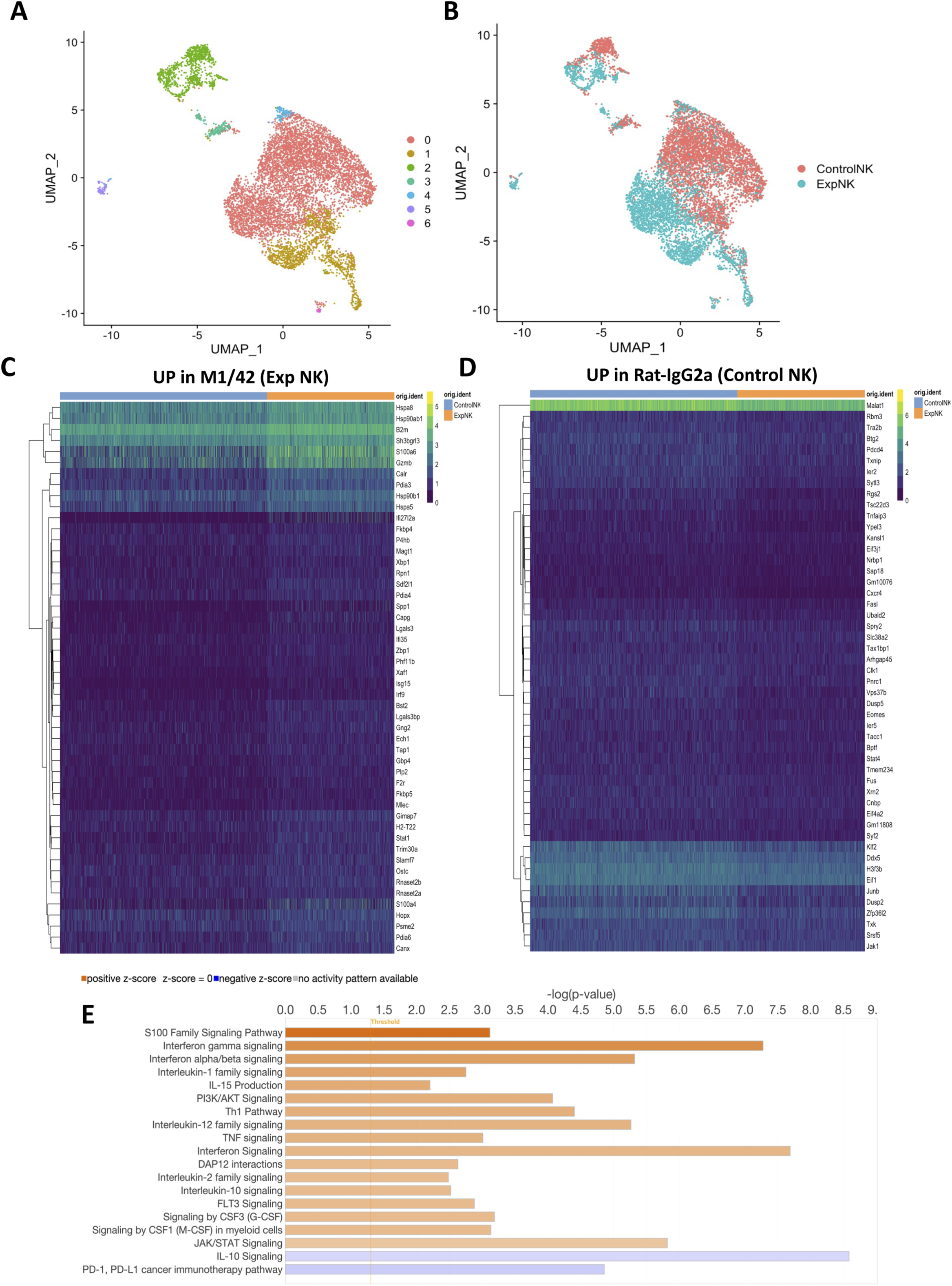
M1/42 treatments increase the expression of Th1 cytokines and interferon-responsive natural killer cell activation genes in established B16F10 metastatic lung tissue. Ten thousand NK1.1^+^ NK cells (CD3^−^CD19^−^CD11b^−^CD11c^−^F4/80^−^NK1.1^+^) were isolated by FACS from the lungs of C57BL/6 mice bearing B16F10 melanoma on day 10 following treatment with either Rat-IgG2a or M1/42 for single cell RNA sequencing. (A) UMAP (Uniform Manifold Approximation and Projection) analysis identified distinct NK cell clusters in both groups; (B) A total of seven clusters (0–6) were observed in treated lung samples. (C) Heat map analysis of gene expression of lung NK cells after M1/42 treatment. (D) Heat map analysis of gene expression of lung NK cells after control rat-IgG2a treatment. (E) Ingenuity pathway analysis based on differential gene expression in metastatic lung-derived NK cells after M1/42 treatment.

### Binding specificity and structural definition of M1/42 interaction with H2 molecules

M1/42 is a pan rat anti-mouse MHC-I mAb that has been used extensively for biological and biochemical studies ^33, 34^. To examine its specificity, we evaluated the binding of M1/42 to a selection of recombinant MHC-I molecules complexed with either human or mouse β_2_m (figure 7A) and determined its affinity (*K*_D_) for H2-K^b^ and H2-D^d^ by surface plasmon resonance (SPR) as 1.9 to 3.8 x 10^−8^ M (figure 7B, C). Consistent with earlier findings, M1/42 binds a variety of murine MHC-I molecules and strongly favors the mouse light chain, β_2_m. We previously reported that M1/42 blocks the binding of soluble recombinant Ly49A and Ly49C proteins to BALB/c and C57BL/6 splenocytes, respectively ^23^. To identify precisely the MHC-I epitope of M1/42, we determined the structure of an M1/42/H2-D^d^ complex by both cryo-EM and X-ray crystallography. The X-ray and cryo-EM models are essentially identical, and since the cryo-EM data refined to higher resolution (2.67 Å vs. 3.60 Å), we confine our description to that model. Data collection and refinement statistics for both structures are presented and compared in online supplemental tables S1 & S2, and details of cryo-EM image classification, experimental protocol, and resolution are shown in online supplemental figure S8A-E. The cryo-EM map and the overall structure of the complex are shown in figure 7D and E, revealing buried surface area of ∼1,000 Å^2^. figure 7F displays the interacting surfaces of H2-D^d^ and M1/42. The details of the interactions (see online supplemental figure S8F, G) include residues of both the H2-D^d^ platform domain (primarily α1 and the first residues of α2) and the β_2_m subunit. The M1/42 H chain contacts 7 residues of the H2-D^d^ H chain (the R14 to F17 loop as well as G90 to S92) and 12 residues of β_2_m (Q2, H34 and E36 and an extensive surface formed by the R81 to P90 loop). The M1/42 L chain contacts residues G16 to E19 and A89. Additional details of the contacts are listed in online supplemental figure S8G. Structure-based amino acid sequence alignment of a selection of murine and human MHC-I molecules with particular attention to the regions of M1/42 contact (residues 14 to 19 and 89 to 92) suggests that the mouse H2 residue 17 (F in H2-D^d^, where human HLA has conserved R) plays a critical role in recognition by M1/42 (online supplemental figure S8H). In addition, β_2_m differences at residues 34, 83, and 85-89 contribute to M1/42 specificity (see online supplemental figure S8I). The refined model of the X-ray crystal structure of the complex at 3.6 Å resolution (8TQ4) is shown in figure 7H (Table S2). The RMSD between X-ray and cryo-EM structures is about 1.3 Å (figure 7I).

**Figure 7.**
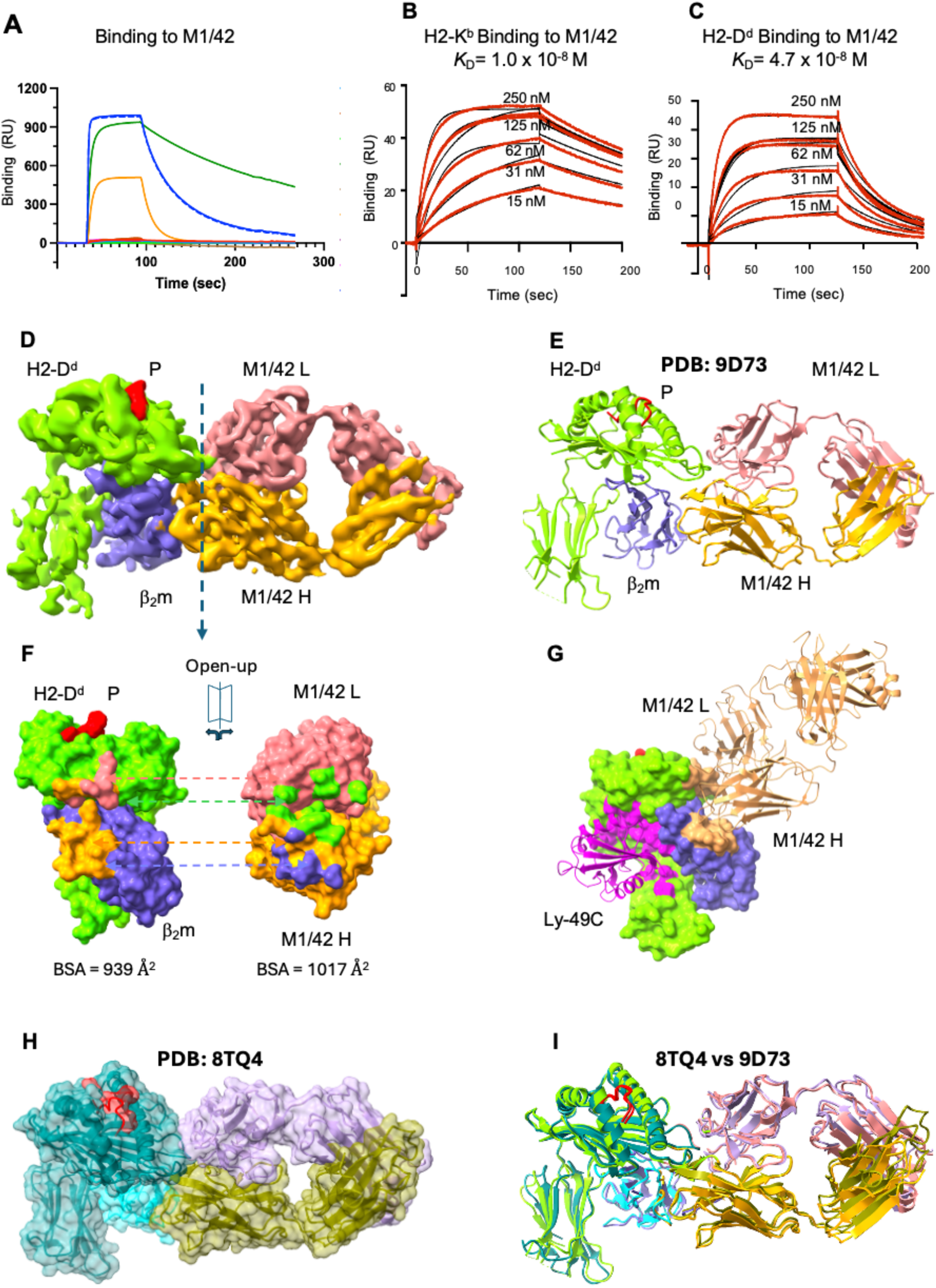
Binding specificity, affinity, and structural definition of M1/42 interaction with H2 molecules. (A) Binding of a panel of human and mouse MHC-I molecules, complexed with either murine or human β_2_m to M1/42. M1/42 was coupled to a BIAcore CM-5 surface as described in Methods and the indicated recombinant proteins were offered in solution phase. Buffer washout was initiated at 100 s. H2-D^d^/mβ_2_m complexes were offered in the first and final injections (solid blue and dashed blue lines) indicating no deterioration of the surface. Other complexes tested, revealing binding not clearly detectable in this display, included: HLA-A*02:01/mouse or human β_2_m; H2-K^b^, H2-D^b^, and H2-D^d^ complexes prepared with their cognate peptides and human β_2_m, as well as mβ_2_m and hβ_2_m alone. (Peptides used in generating the complexes were: for H2-D^d^, RGPGRAFVTI (HIV envelope 318-327); for H2-K^b^, SIINFEKL (chicken ovalbumin 257-264); for H2-D^b^, ASNENMETM (Influenza NP, 366-374); for HLA-A*02:01, GILGFVFTL (influenza M1, 58-66)). (B, C) SPR kinetics binding curves indicating *K*_D_ of interaction of H2-K^b^ or H2-D^d^ for M1/42. Doubling concentrations of the indicated H2 molecules from 15 nM to 250 nM were offered. Binding experiments represent mean ± SEM of three replicates. Kinetics values of *k_a_* for H2-K^b^ and H2-D^d^ were (2.27 ± 0.1) x 10^5^ mol^−1^sec^−1^ and (5.03 ± 0.9) x 10^5^ mol^−1^sec^−1^ respectively; and of *k_d_* of 0.004 ± 0.00003 sec^−1^ and 0.017 ± 0.001 sec^−1^. (D) Cryo-EM density map of M1/42/H2-D^d^/mβ_2_m complex (Electron microscopy data bank (EMDB)-46600, contoured at 0.946); (E) Ribbon diagram of M1/42/H2-D^d^ model (PDB 9D72). (F) Footprints of M1/42 on H2-D^d^, left (surface bound by M1/42 L chain, light coral; surface bound by M1/42 H chain, orange); footprints of H2-D^d^ on M1/42, right (surface bound by H2-D^d^ H chain, chartreuse; surface bound by β_2_m, slate blue). (G) Footprints of M1/42 and Ly49C on H2 do not overlap. M1/42/H2-D^d^ (PDB 9D72) and Ly49C/H2-K^b^ (PDB 3C8K) were superposed based on the H2 H chains and contact surfaces to M1/42 and Ly49C were visualized in ChimeraX ^45^. Ly49C lectin domain is magenta, M1/42 H and L chains are sandy brown. H) Surface representation of X-ray crystal structure of M1/42/H2-D^d^/mb_2_m. (I) Superimposed X-ray (8TQ4) and cryo-EM (9D72) structures of M1/42/H2-D^d^/mβ_2_m, RMSD = 1.3 Å.

## DISCUSSION

The findings presented in this study highlight the remarkable potential of antibody-mediated blockade of innate receptor Ly49/MHC-I interactions as a strategy to unleash both innate and adaptive immunity against cancer metastasis. The pan anti-MHC-I monoclonal antibody M1/42 demonstrates a unique mechanism of action, distinguishing itself from conventional checkpoint inhibitors by targeting MHC-I rather than individual inhibitory receptors on natural killer (NK) cells. This approach simultaneously interferes with multiple inhibitory pathways, leading to broad activation of immune responses, including NK cells, memory T cells, and myeloid cell subsets.

One of the key observations is that M1/42 administration robustly stimulates NK cell proliferation and activation independent of T cells and Fcγ receptors, as evidenced by experiments in Rag^−/−^ and FcγR^−/−^mice. The antibody’s ability to enhance proliferation and activation of NK cells, memory CD4 and CD8 T cells, dendritic cells, and macrophages in both lymphoid (spleen) and non-lymphoid (liver) tissues underscores its systemic immunostimulatory effects. Notably, the expansion of effector and central memory T cells, as well as the upregulation of Th1-type transcription factors (TCF-1, T-bet), points to a coordinated activation of innate and adaptive immunity.

The therapeutic efficacy of M1/42 is particularly striking in models of checkpoint inhibitor-resistant cancers, such as PDAC and B16F10 melanoma. In the PDAC models harboring KRASG12D and TP53R172H mutations, which are notoriously refractory to anti-CTLA4 and anti-PD1 therapies ^31, 35, 36^, M1/42 treatment significantly restricted both primary tumor growth and metastatic colonization in the liver and lungs. This was accompanied by an increased frequency of tumor-infiltrating CD4^+^ and CD8^+^ T cells, a reduction in Tregs, and an expansion of dendritic cells at the tumor site.

Similarly, in the B16F10 melanoma lung metastasis model, M1/42 administration led to a marked reduction in metastatic burden, increased infiltration of CD8^+^ T cells, and enhanced expansion of tumor-specific memory CD8^+^ T cells (GP100^+^ and Trp2^+^). Depletion experiments revealed that NK cells are indispensable for the anti-metastatic effects of M1/42, while CD8 T cells play a supportive role ^31^. Compared to classical checkpoint inhibitors, M1/42 more effectively promoted NK cell proliferation, reduced Treg frequencies, and drove the expansion of effector memory T cells, indicating a broader and more potent immunomodulatory effect. The elevation of Th1-type cytokines and chemokines in serum further supports the notion of a pro-inflammatory, anti-tumor microenvironment induced by M1/42.

Mechanistically, single-cell RNA (scRNA) sequencing of metastatic lung tissue following M1/42 treatment revealed upregulation of genes involved in antigen processing and presentation, interferon gamma responsiveness, and Th1 cytokine production. Pathway analyses indicated enhanced signaling through interferon alpha/beta, IL-15, PI3K/Akt, Th1, and FLT3 pathways, alongside downregulation of PD1/PDL1 inhibitory signals. These molecular changes provide insight into how M1/42 orchestrates a multifaceted immune response capable of overcoming tumor-induced immunosuppression and resistance to checkpoint blockade.

Our working model for the functional effects of M1/42 *in vivo* was based on its ability to block *in vitro* the binding of biotinylated, dimeric, full-length Ly49 proteins that included both the globular lectin domain and the long flexible stalk region, to MHC-I on mouse splenocytes ^23^. This initial view was supported by our demonstration of direct competition between the human pan anti-HLA mAbs DX17 and W6/32 for the LILR binding site on HLA ^24^. However, comparison of the H2-D^d^ footprint of M1/42 in both our cryo-EM and X-ray structures reported here with that of the H2-K^b^ footprint of Ly49C (PDB 3C8K) ^37^, shows that M1/42 and Ly49C bind to distinct non-overlapping sites of MHC-I (Fig 7G), indicating that direct competition does not easily explain the functional blocking by the antibody. Instead, the functional competition may result from allosteric effects or steric hindrance due to bivalent engagement of MHC-I by the antibody and the consequent limited accessibility of a full-length Ly49 dimer. The structures of the several crystallographically determined Ly49/MHC complexes (1QO3 ^38^, 1P4L ^11^, 1P1Z ^39^, 3C8K ^37^, 5J6G ^40^) have only revealed the lectin domain and thus the structure of a dimeric, full-length Ly49 remains elusive. Ly49/H2 interactions have been shown to assume either a *cis* (i.e. interaction of Ly49 with H2 molecules on the same cell) or *trans* (interaction between different cells) orientation ^41–43^. It remains possible that M1/42 inhibits the formation of a Ly49/MHC-I complex necessary for signal transduction. Others have offered a similar model for the blocking of some human NK receptors by the human pan anti-HLA molecule W6/32 ^44^. This structural insight could guide the development of new antibodies that broadly target MHC-I interactions, offering therapeutic advantages over receptor-specific blockade.

In summary, M1/42-mediated blockade of MHC-I (H2) and NK-specific Ly49 innate receptor interactions represents a paradigm shift in immunotherapy, opening new avenues for the treatment of metastatic cancers that are refractory to existing checkpoint inhibitors. By harnessing the synergistic potential of innate and adaptive immunity, this approach may pave the way for more effective and durable anti-cancer therapies. Future studies should investigate the long-term effects of MHC-I blockade, potential off-target immune activation, and the applicability of this strategy in other tumor types and in clinical settings by targeting HLA/KIR ^25^ or HLA/LILR ^24^ interactions alone or in combination with classical checkpoint inhibitors or CAR-T cell therapy.

## Supporting information

Panda et al Supplemental Files

## DECLARATIONS

### Ethics approval

This study involved no human experiments, and all animal experiments approved by institutional animal care and use committee.

### Consent for publication

All authors consent to publication of this paper.

### Data availability statement

Data are available as follows: RNA-seq data have been deposited at the Gene Expression Omnibus (GEO) site as GSE329297. The cryo-EM map is available at EMDB-46600, protein data bank (PDB) model 9D72, and the X-ray data and coordinates are available at PDB 8TQ4. Other unique reagents may be obtained from the authors.

### Competing interests

The authors declare no competing interests.

### Funding

This research was supported by the Intramural Research Program of the National Institutes of Health (NIH). The contributions of the NIH author(s) were made as part of their official duties as NIH federal employees, are in compliance with agency policy requirements, and are considered Works of the United States Government. However, the findings and conclusions presented in this paper are those of the author(s) and do not necessarily reflect the views of the NIH or the U.S. Department of Health and Human Services.

### Authors’ contributions

AKP, SSinha, and KN conceived the project, performed experiments, analyzed data, and wrote the paper. JJ performed X-ray and cryo-EM experiments, analysis, and writing. SC, SK, Y-HK, SSharma and PS performed experiments and analyzed data. JMH, DHM, and EMS designed and supervised the project, analyzed data and wrote the paper.

## Acknowledgements

X-ray data were collected at Southeast Regional Collaborative Access Team (SER-CAT) 22-ID or 22-BM beamlines at the Advanced Photon Source, Argonne National Laboratory. SER-CAT is supported by its member institutions (www.ser-cat.org/members.html) and equipment grants (S10_RR25528 and S10_RR028976) from the National Institutes of Health. Use of the Advanced Photon Source was supported by the U.S. Department of Energy, Office of Science, Office of Basic Energy Sciences, under Contract No. W-31-109-Eng-38. The Electron Microscopy Resource is supported by the National Cancer Institute and NIH Intramural Research Program Cryo-EM Consortium (NICE). We thank Drs. Haotian Lei (RTB, NIAID) and Rick Huang (NICE, NCI) for their help in grid screening and cryo-EM data collection.

## ABBREVIATIONS

PDAC: pancreatic ductal adenocarcinoma
cryo-EM: cryo-electron microscopic analysis
EMDB: electron microscopy data bank
PDB: protein data bank
NK: natural killer
MHC-I: major histocompatibility complex class I molecule
KIR: killer cell immunoglobulin-like receptor
LILR: leukocyte immunoglobulin-like receptor
MP: memory phenotype
FcR: Fc receptor
KPC (LSL-Kras G12D/+; LSL-Trp53 R172H/+; Pdx-1-Cre): pancreatic cancer mouse model
FBS: fetal bovine serum

## Notes

### Competing Interest Statement

The authors have declared no competing interest.

