## Supplementary material for "Antibody Blockade of Ly49/MHC-I interactions enhances Innate and Adaptive Immunity Against Cancer Metastasis": Panda et al Supplemental Files

**SUPPLEMENTAL MATERIALS**

Online Supplemental Materials and Methods

Online Supplemental Figure Legends

Online Supplemental Tables

Online Supplemental Figures

### ONLINE MATERIALS AND METHODS

#### Animals

C57BL/6 mice were purchased from Charles River Laboratories. $Fc\gamma R^{-/-}$ mice obtained from Taconic Biosciences. $\beta_2m^{-/-}$ mice were procured from Jackson Laboratories. All mice were sex- and age-matched for experimentation and used between 8 and 12 weeks of age. The KPC mice (LSL-Kras G12D/+ ;LSL-Trp53 R172H/+ ;Pdx-1-Cre ) were procured from NCI inventory at Frederick. All animal protocols used in this study were approved by the National Institute of Allergy and Infectious Diseases Animal Care and Use Committee (LISB-14 and LISB-15E).

#### Cell Lines

The B16F10 cell line was purchased from ATCC and the syngeneic mouse KPC cell was provided by Dr. Serguei Kozlov at the NCI**.** All cell lines were mycoplasma-tested by PCR and maintained in complete DMEM supplemented with 10% FBS, 1× penicillin/streptomycin, and 2.5 µg/mL Plasmocin for eliminating and preventing mycoplasma-related effects.

#### Splenocyte isolation

Spleens were harvested from naive and tumor bearing mice on indicated days and the tissues were homogenized and cells passed through a 0.70-µm cell strainer. Red blood cells were lysed using sterile ACK lysing buffer [$NH_4Cl$ (0.15 M), $KHCO_3$ (10 mM) and $Na_2EDTA$ (0.1 mM), pH 7.3]. Lymphocytes were washed, suspended in sterile complete medium [RPMI medium supplemented with 10% heat-inactivated fetal bovine serum (FBS), L-glutamine (2 mM), sodium pyruvate (1 mM), HEPES (1 mM), non-essential amino acids (0.1 mM), 2-mercaptoethanol (50 μM), and penicillin and streptomycin (100 U/ml)], and total live cells were counted.

**Isolation of Liver and Lung Lymphocytes**

Livers and lungs were harvested from euthanized mice and mechanically dissociated using a 0.75-μm cell strainer and the rubber end of a 3cc syringe into a 50 mL tube containing 5-10 mL RPMI medium per organ. The resulting cell suspension was centrifuged at 300 x g for 3 minutes to pellet hepatocytes and tumor cells; the supernatant was then transferred to a new tube. After centrifugation at 1,500 rpm for 5 minutes, the cell pellet was resuspended in 32% Percoll™ solution and subjected to centrifugation at 2,000 rpm, 14°C with no brake for 15 minutes. Red blood cells were lysed by washing the cell pellet with 5 mL ACK lysis buffer, followed by centrifugation at 1,500 rpm for 5 minutes. The purified immune cell pellet was washed twice in FACS staining buffer [PBS supplemented with 10% heat-inactivated fetal bovine serum (FBS)] and subsequently used for surface and intracellular antibody staining.

**Flow cytometry**

Mouse lymphocytes were prepared as single-cell suspensions (≈0.5×10^6 cells) from spleen or tumors from liver and lungs and resuspended in staining buffer (PBS, 10% heat-inactivated FBS, 0.05% sodium azide). Cells were Fc-blocked (TruStain) for 5 min, stained with a viability dye and surface antibody cocktails for 30 min at 4°C, washed, and, where indicated, incubated with MHC-I tetramers from the NIH Tetramer Core Facility (H2-$D^b$ GP100, H2$K^b$ Trp-2) at 1:50 for 1 h at

4°C. For intracellular transcription factor and proliferation analysis, surface-stained cells were fixed and permeabilized using a Foxp3 transcription factor buffer set and stained for Foxp3, Ki-67, Tbet, & Eomes. For functional assays, cells were stimulated in complete RPMI with PMA/ionomycin plus brefeldin A and monensin for 4 h with GolgiStop, then surface-stained, fixed/permeabilized, and stained for intracellular cytokines (IFN-γ, TNF-α, Granzyme B). Samples were acquired on a BD LSRFortessa and analyzed using FlowJo v10.

**Antibody Treatment**

Rat-IgG2a (250μg/dose, i.p., Clone 2A3, BioXcell) and pan anti-MHC-I (H2) (500μg/dose, Clone M1/42, BioXcell), anti-CTLA4 (250μg/dose, Clone UC10-4F10-11, BioXcell), anti-PD1 (Clone 29F.1A12, BioXcell) anti-PDL1 (250μg/dose, Clone 10F.9G2, BioXcell) or anti-NKG2A (250μg/dose, Clone 20D5, BioXcell) were given to naïve or tumor bearing mice as indicated in the figure legends. For NK or CD8 T cell depletion, 400ug/dose anti-NK1.1 (clone: PK136, BioXcell) or anti-CD4 (Clone: GK1.5, BioXcell) or anti-CD8 (Clone: 2.43, BioXcell) were given intra-peritoneally (i.p) on two successive days followed by M1/42 co-treatment every 2-3 days starting from day 0 and ending on day 6 or day 15.

**Immunohistochemistry**

B16F10 tumor bearing Lungs on day 10 were perfused with 1X PBS to remove blood and tracheally inflated with 10% neutral buffered formalin (NBF) using a 20G needle. Excised whole lungs were submerged in 10% NBF for 24 hours at 4°C and then transferred to 70% ethanol. Following dehydration in graded alcohols and clearing with xylene, lungs were embedded in paraffin wax. Sections (4mm) were cut using a microtome and mounted on charged slides. Slides were then deparaffinized, rehydrated, and subjected to antigen retrieval and blocking according to established protocols. Primary antibody incubation was performed overnight at 4°C, followed by HRP-conjugated secondary antibody for 1 hour at room temperature. Detection was performed with DAB chromogen and counterstained with Hematoxylin. Stained slides were scanned at 20X magnification using a high-resolution Leica Aperio scanner and analyzed with ImageScope software.

**Cytokine Bead Array**

Mouse whole blood was collected in a heparinized tube. After centrifugation at 2000 x g for 15 min, plasma was transferred to a new tube. Concentrations of cytokines were determined by LEGENDplex™ Cytometric Bead Array (Biolegend) following the manufacturer's instructions.

**Single-cell RNA-seq library preparation and sequencing**

NK cells ($CD3^{-}CD19^{-}CD11b^{-}CD11c^{-}NK1.1^{+}$) were sorted from lung lymphocytes of B16F10 melanoma–bearing C57BL/6 mice treated with Rat-IgG2a or M1/42 on days 3, 6, and 9. Lungs were harvested on day 10. Single-cell RNA-sequencing libraries were prepared using the Chromium Next GEM Single Cell 3' Reagent Kits v3 (10X Genomics) and processed on the Chromium System (10X Genomics), according to manufacturer's protocol. Briefly, single cell suspensions were loaded onto the Chromium Next GEM Chip B microfluidics chip at a target recovery of 10,000 cells per sample, along with prepared master mix, 10X Barcoded gelbeads, and partitioning oil. Single cells were partitioned into Gelbead-in-Emulsion (GEM) droplets, where 10X Barcoded gelbeads captured the poly-adenylated transcriptome of each cell and proceeded through reverse transcription, resulting in full-length cDNA uniquely barcoded per individual cell. Dual-Indexed libraries were then generated using 13 cycles of PCR amplification, following manufacturer's instructions.

Final libraries were pooled in equimolar ratios and sequenced on a NextSeq 500 Sequencing System (Illumina), using NextSeq 500 High Output (150 cycle) reagent kits (Illumina). Sequencing depth was targeted to achieve 50,000 reads per cell, using the following cycles of chemistry as per 10X Genomics recommended sequencing parameters: Read1=28 cycles; Index1=10 cycles; Index2=10 cycles; Read2=90 cycles. GEX libraries were barcode processed and the sequencing reads were aligned against mm9 reference genome and gene counting were done using 10X Genomics Cell Ranger v6.1.2

**Pathway Analysis**

Gene-level quantification was performed with RSEM v1.3.1 and differential expression testing with DESeq2 (FDR < 0.05). Pathway enrichment was assessed by GSEA using GO and KEGG gene sets, and Reactome analysis of B16F10-bearing NK cell DEGs was performed via STRING. DEGs from B16F10-derived NK cells were analyzed in Ingenuity Pathway Analysis (IPA; Qiagen); canonical pathway significance was determined by Fisher's exact test ($P < 0.05$, $-\log_{10}P > 1.3$), activation/inhibition by IPA Z-score ($Z > 2$ = activated; $Z < -2$ = inhibited), and pathways were ranked by Z-score (activated = positive/orange; inhibited = negative/blue).

**Liver Metastasis, Splenic Injection and Liver Outgrowth Assay**

Liver metastatic outgrowth and colonization were evaluated through intrasplenic injections. Briefly, the mouse is anesthetized, and an injection of 2 mg/kg of meloxicam and 0.5 mg/kg of buprenorphine is administered subcutaneously. Sterile eye lubricant is applied to both eyes to prevent corneal drying during surgery. When administering injections into the C57/BL6 mouse, the fur covering the abdominal wall is carefully removed by applying hair removal cream. The abdominal skin is then cleaned with a moist gauze pad to remove loose hair and sterilized with

Betadine and 70% ethanol. A small volume of local anesthetic agent, such as bupivacaine (0.25%-0.5%), is injected into the tissue adjacent to the intended incision line. A 0.5- to 1-cm parasagittal incision is made in the skin of the left flank using sterile scissors, revealing the abdominal muscles. Sterile scissors are then used to make a 0.5- to 1-cm parasagittal incision through the abdominal musculature over the spleen. The spleen is exteriorized through the incision by gentle retraction with forceps and held in place using a sterile cotton–tipped applicator or placed on a sterile moist gauze pad. A total of 0.3 x $10^6$ tumor cells in 100 μL of 1xPBS are injected into the splenic parenchyma under the splenic capsule using a 26-gauge needle. When the needle is withdrawn, gentle pressure is applied to the injection site using a sterile cotton swab or gauze for 3 to 5 minutes or until hemostasis is achieved. Three to five minutes after splenic injection when adequate hemostasis is achieved, the spleen is removed, and the splenic artery and vein are ligated using 6-0 suture. The abdomen is checked for bleeding. If bleeding is found, gentle pressure is applied using a cotton swab or sterile gauze until hemostasis is achieved. The muscle layer is closed using sterile absorbable suture (e.g., Vicryl) of the appropriate diameter in a simple interrupted pattern. Skin edges are opposed and closed with a monofilament absorbable suture (e.g. Monocryl) of the appropriate diameter in a subcuticular pattern. Immediately following surgery, when the mouse is under anesthesia, 100 μL of luciferin is given by retroorbital injection to visualize tumor cells in the liver via IVIS imaging. Thereafter, mice were imaged weekly or at an experimental endpoint to visualize tumor outgrowth and colonization in the liver.

**Tail-vein injection for lung metastasis**

Mice were warmed under a heat lamp for 5-10 minutes to promote dilation of the tail veins. After warming, each mouse was secured in a standard restrainer (Braintree Scientific) to ensure stable access to the tail vein. Once the vein was identified, 0.3 x $10^6$ tumor cells (KPC or B16F10) suspended in 100 µL of 1 x PBS were carefully injected using a 28-gauge needle, taking care to minimize the risk of vascular overload or rupture. Following injection, gentle pressure was applied to the site with sterile gauze for approximately 30 seconds to prevent hematoma formation. Immediately after the procedure, mice were anesthetized and received 100 µL of luciferin via retro-orbital injection, enabling *in vivo* visualization of tumor cells within the lungs using IVIS imaging. Mice were subsequently imaged at weekly intervals or at the designated experimental endpoint to monitor tumor outgrowth and lung colonization.

**IVIS imaging and metastatic outgrowth analysis**

To assess metastatic outgrowth and the colonization capacity of tumor cells, recipient mice were anesthetized with isoflurane and injected retro-orbitally with 100 µL of luciferin (15 µg/mL). Whole-body bioluminescent images were acquired using the IVIS Spectrum system (PerkinElmer) at specified timepoints. For liver colonization studies in C57BL/6 mice, the abdominal fur was removed using hair removal cream prior to luciferin administration and imaging; the liver region was defined as the area between the sternum and the middle of the abdomen. For endpoint analysis of whole organs (liver or lung), mice were injected with luciferin, humanely euthanized, and their livers and lungs were excised for *ex vivo* imaging on the IVIS Spectrum. Quantification of liver and lung colonization was performed using Living Image software (versions 4.5.2 and 4.7.2), by drawing regions of interest (ROIs) over the defined anatomical areas and measuring the bioluminescent signal in radiance (photons/sec/cm2/sr). The software automatically normalizes for sensitivity differences due to variable exposure times, ensuring that ROI values are reported as calibrated physical units without requiring manual adjustment.

**Cryo-EM sample preparation and data collection, structure determination and refinement**

Freshly purified M1/42 Fab was mixed in a 1:1 molar ratio with bacterially expressed and refolded H-2D$^d$ prepared as described previously [1] and the complexes were purified by size exclusion chromatography (SEC). Complexes at a concentration of 0.7-1.4 mg/ml were applied onto holey-carbon cryo-EM grids (C-flat TM Holey Carbon Grid Gold 1.2 μm/1.3 μm space 300 mesh (Protochip, NC USA)), which had been glow discharged for 60 seconds. Grids were blotted for 3 seconds, and plunged into liquid ethane with a Vitrobot Mark4 (Thermo-Fisher). Cryo-EM data were collected on a Titan Krios 300-keV microscope (NICE/NIH Cryo-EM consortium). Images were acquired automatically with SerialEM 76 on a BioQuantum-K3 detector (Gatan) in super-resolution mode at 130x nominal magnification (0.83 Å/unbinned pixels) and a defocus range from -0.7 to -2.0 μm. An exposure time of 0.05s per frame was used, with a total exposure of about 54.2 electrons/Å$^2$. 2,993 movies were recorded. Image processing, 2D classification, 3D reconstruction, and map refinements were performed using cryoSPARC v4.4.1 [2] following their standard procedures. To improve the map resolution for the relatively small Fab/MHC-I complex (MW ~90 kDa), we applied a recently developed protocol for particle picking [3], and we were able to extend the high resolution from 4.0Å to 2.67Å (Fig. S8). The total particles selected at each step are listed in Fig.S8C.

We used the X-ray crystal structure of the M1/42 Fab/H-2D$^d$ complex (PDB: 8TQ4) to dock and manually fit the cryo-EM maps of Fab M1/42+H2-D$^d$. Real space refinement was carried out in Phenix [4] which included rigid body refinement, simulated annealing at the initial step, local grid search, ADP refinement, and the application of secondary structure restraints. The final refined model compared with the map density has an overall Correlation Coefficient (CC) of 0.75/0.74/0.68 (mask/volume/peaks) for Fab M1/42+H2-D$^d$. We also calculated the Q-score for individual residues [5] for validation. Cryo-EM Data processing, refinement statistics, and model validation are listed in online supplemental table S1.

**Protein Expression, purification, crystallization, data collection, and X-ray structure determination**

M1/42 was purified from hybridoma culture supernatant by immunoadsorbent chromatography on Protein G-Sepharose and Fab fragments were prepared as previously described for other antibodies [3]. Recombinant H2-$D^d$ (extracellular domains) complexed with P18-I10 (HIV IIIB envelope glycoprotein peptide 318-327, RGPGRAFVTI) and murine β2m was made as described[6]. Crystallization conditions were identified by screening hanging drops of M1/42+H2-$D^d$ at 18 °C. Crystals of M1/42+H2-$D^d$ were grown in 10% PEG 4000, 0.1M Sodium citrate pH 5.5, 0.2M Sodium acetate, cryoprotected in mother liquor containing 10% ethylene glycol, and flash frozen in liquid nitrogen. Diffraction data were collected (at wavelength 1.033 Å, in $N_2$ stream at ~ 100 K) at Southeast Regional Collaborative Access Team (SER-CAT) beamline 22ID at the Advanced Photon Source, Argonne National Laboratory and processed with XDS [7] to 3.6 Å resolution for M1/42+H2-$D^d$ (**see Table.1**). The structures were solved by molecular replacement with Phaser [8] using H2-$D^d$ from PDB 5WEU as the search model. We used the model of DX17 Fab (8TQ6) with the CDR loops trimed off as the initial search model for M1/42, then manually rebuilt the CDR loops according to the electron densities. These molecular replacement models were subjected to several rounds of refinement with Phenix [4] interspersed with manual building in Coot [9]. Fab sequences were determined by PCR sequencing. $R_{work}/R_{free}$ (%) values for final refined models of M1/42+H2-$D^d$ are 22.1/26.0. Data collection and refinement statistics are summarized in online supplemental table S1. Graphics figures were generated with PyMOL.

**Surface Plasmon Resonance Experiments**

The kinetics of the interaction of M1/42 with MHC-I molecules were examined by SPR in a BiaCore T200 (Cytiva) at 25°C in 10 mmol/L TRIS pH 7.4, 150 mmol/L NaCl, 3 mmol/L EDTA, and 0.05% surfactant P20. Antibody was immobilized on the surface of a CM5 chip by EDS/NHC coupling chemistry and graded concentrations of MHC-I were sequentially injected over the M1/42 surface at a flow rate of 30 μL/min. Regeneration of the surface at the end of each binding and washout cycle was with 0.1M glycine pH 2.5. In binding experiments with H-2$D^d$ and H-2$K^b$, M1/42 was captured on immobilized goat anti-rat IgG antibody. Binding experiments were repeated three times. Sensorgrams were fit globally to a 1:1 binding model with BiaCore T200 Evaluation Software 3.1 and plotted with Prism (GraphPad Software).

### Statistical analysis

Statistical comparisons were made by two-tailed unpaired t test or one-way ANOVA (Prism 10); $P < 0.05$ was considered significant.

### SUPPLEMENTAL FIGURE LEGENDS

**Figure S1** C57BL/6 and Fcγ$R^{-/-}$ mice treated with antibodies every other day for 6 days and splenocytes were harvested on day 8. (A) C57BL/6 animals treated with M1/42 for six days and analyzed for proliferation (Ki-67 incorporation) of NK cells, $CD4^+CD44^+CD62L^-$, $CD8^+CD44^+CD62L^+$, and $CD8^+CD44^+CD62L^-$. (B-E) Fcγ$R^{-/-}$ mice treated with M1/42 and splenocyte-derived NK, CD4 and CD8 memory were analyzed for cell proliferation on day 8.

**Figure S2** (A & B) KPC cells ($2x10^5$) were injected (s.c.) into the right flank of C57BL/6 animals, M1/42 was administered every 3-4 days from day 3 to day 20, and tumor volume was compared on day 22. (C) KPC-Luc2 cells were injected into the spleen, followed by splenectomy at 5 minutes. Luciferase signals were measured by *in vivo* bioluminescence imaging every week to determine metastatic outgrowth and colonization in the after Rat-IgG2a or M1/42 treatment on day 0, day 14 and day 21. (D) *In vivo* bioluminescence quantification of photon flux of liver metastatic outgrowth on day 0, 14, and 21 after Rat-IgG2a and M1/42 treatment. (E ) KPC cell outgrowth on metastatic liver was measured by bioluminescence imaging on day 22 after Rat-IgG2a and M1/42 treatment (5 mice per group) every 3-day interval starting from day 14. (F) Quantification of luciferase signals (photons/second) of the harvested lungs of each group as represented by a graph. (G) Flowcytometric and statistical analysis of TILs derived from liver on day 22 after antibody treatments. Percentage of CD4 and CD8 T cells, (H) $CD4^+FOXP3^+$ Treg, (I) $CD4^+Foxp3^-CD44^+CD62L^-$,(J) $CD8^+CD44^+$ T cells and (K) $CD19^-CD11c^+$ dendritic cells.

**Figure S3** KPC-Luc-2 cells were injected intravenously (tail vein) into C57BL/6 mice to induce lung metastasis. From day 14 to 28, animals received rat-IgG2a or M1/42 antibody every three days, and lungs were collected on day 30. Representative flow cytometric plot and statistical analysis of (A) Percentage CD4 and CD8 T cells, (B) Percentage of $CD4^+Foxp3^+$ Treg cells (C) Percentage of $CD4^+Foxp3^-$ effector T cells, (D) Percentage of $CD4^+Foxp3^-CD44^+CD62L^-$ effector Memory T cells proliferation (Ki-67), (E) Percentage of memory CD8 T cell subsets, (F) Percentage of Ki-67 expression on memory $CD8^+CD44^+$ T cell subsets (G) Percentage of $CD19^+$ B cells and $NK1.1^+$ NK cells (H) Relative Ki-67 expression in NK cells (I) Percentage of $CD11b^+Gr1^+$ monocytes.

**Figure S4** (A) C57BL/6 mice ($1.5 \times 10^5$ cells) were injected in the tail vein with B16F10 melanoma cells on day 0. Subsequently, rat-IgG2a and M1/42 were administered every 3–4 days from day 3 to day 18. (B) Metastatic lung outgrowth was analyzed on day 20 following treatment with rat-IgG2a or M1/42. (C) CD8 immunohistochemistry was performed on B16F10 lung samples obtained on day 15 after administration of rat-IgG2a and M1/42 on days 3, 6, 9, and 12. (D-F) C57BL/6 mice received intravenous B16F10 melanoma cell injections on day 0, followed by scheduled administration of rat-IgG2a, M1/42, anti-CTLA4, anti-PD1, or anti-PDL1 in combination with anti-NKG2A every 3–4 days from day 3 to day 20. (D) Relative percentage of $CD4^{+}Foxp3^{+}$ Tregs of metastatic lung lymphocytes. (E) Percentage of $CD4^{+}CD44^{+}Foxp3^{-}CD44^{+}$ T cells. (F) Percentage of $CD8^{+}CD44^{+}$ T cells in metastatic lungs in antibody-treated groups on Day 22. (G & H) Flowcytometric contour plot depicting the percentage of $IFN\gamma^{+}$Granzyme-$B^{+}GP100^{+}$ and $IFN\gamma^{+}$Granzyme-$B^{+}TRP2^{+}$ melanoma-specific $CD8^{+}$ T cells in metastatic lungs, determined on day 22 after M1/42 and checkpoint inhibitor treatment.

**Figure S5** Ten thousand $NK1.1^{+}$ natural killer cells ($CD3^{-}CD19^{-}CD11b^{-}CD11c^{-}F4/80^{-}NK1.1^{+}$) were isolated by FACS from the lungs of C57BL/6 mice bearing B16F10 melanoma on day 10 following treatment with either rat-IgG2a or M1/42 for bulk RNA sequencing. (A) Reactome pathway analysis of upregulated gene expression (top 50 genes) after M1/42 treatment as compared with isotype control group. (B) Reactome pathway analysis of down-regulated gene expression (top 50 genes) after M1/42 treatment as compared with isotype control group. (C) KEGG pathway analysis of differential gene expression (top 54 genes) after M1/42 treatment as compared with isotype control group. (C) GO pathway analysis of differential gene expression (top 54 genes) after M1/42 treatment as compared with isotype control group.

**Figure S6** Heat map analysis demonstrated regulation of gene expression in NK cells after M1/42 treatment in lungs in (A & B) cluster 0, (C & D) cluster 1 and (E & F) cluster 2.

**Figure S7** Reactome pathway analysis of upregulation (top 50 genes) and down-regulation (top 50 genes) of gene expression in NK cells after M1/42 treatment in lungs in (A & B) cluster 0, (C & D) cluster 1 and (E & F) cluster 2.

**Figure S8** M1/42 + H2-$D^d$ cryo-EM: 2D classification and protocol; interactions and contacts. **(**A) 2D classification of cryo-EM images after several runs of particle picking. (B) The best 2D class shows domains of Fab and MHC-I. (C) The protocol used in template picking and the number of resulting particles in each step. (D) Map resolution as indicated by Fourier Shell Correlation (FSC) at 0.143 reveals 2.67 Å resolution (no mask = 3.6 Å). (E) Local resolution visualization on the surface of M1/42+H2-$D^d$. Blue is higher resolution, red is lower resolution. (F) M1/42 contacts between Fab H chain and L chain and H2-$D^d$ H chain and $\beta_2$m for PDB 9D72 were tabulated from PDBsum [10]; (G) Residue contacts and distances taken from PISA analysis [11] are summarized; (H) MHC variability for a sampling of HLA or H2 class I sequences was performed for the indicated amino acid residues for HLAs A*01:01:01:01, A*11:01:01:01, A*02:01:01:01, A*03:01:01:01, B*13:01:01:01, B*35:01:01:01, B*27:05:02:01, C*05:01:01:01, C*06:02:01:01, E*01:01:01:01:01, F*01:01:01:01, and G*01:01:01:01 (notation according to IMGT database [12]); and an alignment of amino acid sequences of H2 from UniProt: $D^d$ (P01900), $K^d$ (P01902), $L^d$ (P01897), $K^b$ (PO1901), $D^b$ (P01899), $K^k$ (P04223), and $D^k$ (P14426). Sequences were aligned with ClustalW as implemented in MacVector 18.18.2 and residues 14 to 19 were analyzed and for HLA displayed with WebLogo [13] (https://weblogo.threeplusone.com/create.cgi). (I) $\beta_2$m polymorphism at contacts.

**Fig. S1**

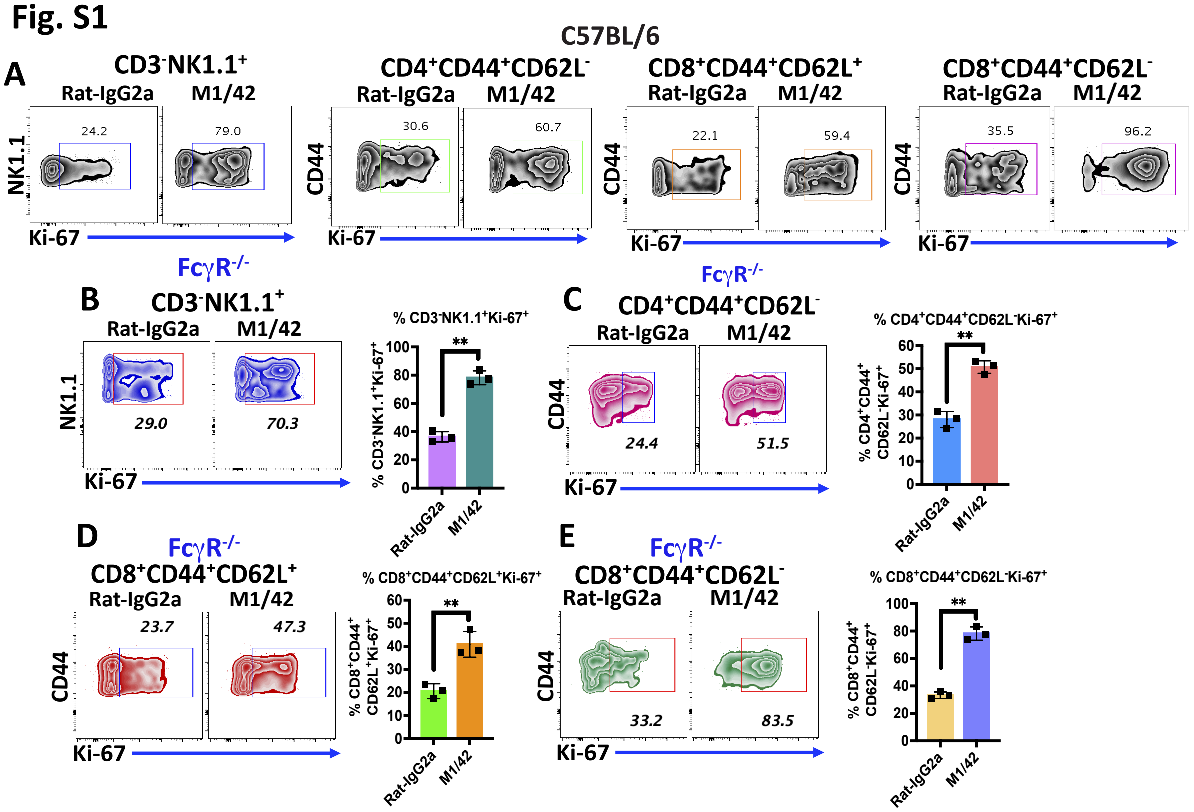
C57BL/6
A
CD3⁻NK1.1⁺
Rat-IgG2a
M1/42
24.2
79.0
NK1.1
Ki-67
CD4⁺CD44⁺CD62L⁻
30.6
60.7
CD44
CD8⁺CD44⁺CD62L⁺
22.1
59.4
CD8⁺CD44⁺CD62L⁻
35.5
96.2
FcγR-/-
B
CD3⁻NK1.1⁺
29.0
70.3
% CD3⁻NK1.1⁺Ki-67⁺
**
C
CD4⁺CD44⁺CD62L⁻
24.4
51.5
% CD4⁺CD44⁺CD62L⁻Ki-67⁺
D
CD8⁺CD44⁺CD62L⁺
23.7
47.3
% CD8⁺CD44⁺CD62L⁺Ki-67⁺
E
CD8⁺CD44⁺CD62L⁻
33.2
83.5
% CD8⁺CD44⁺CD62L⁻Ki-67⁺

**Fig-S2**

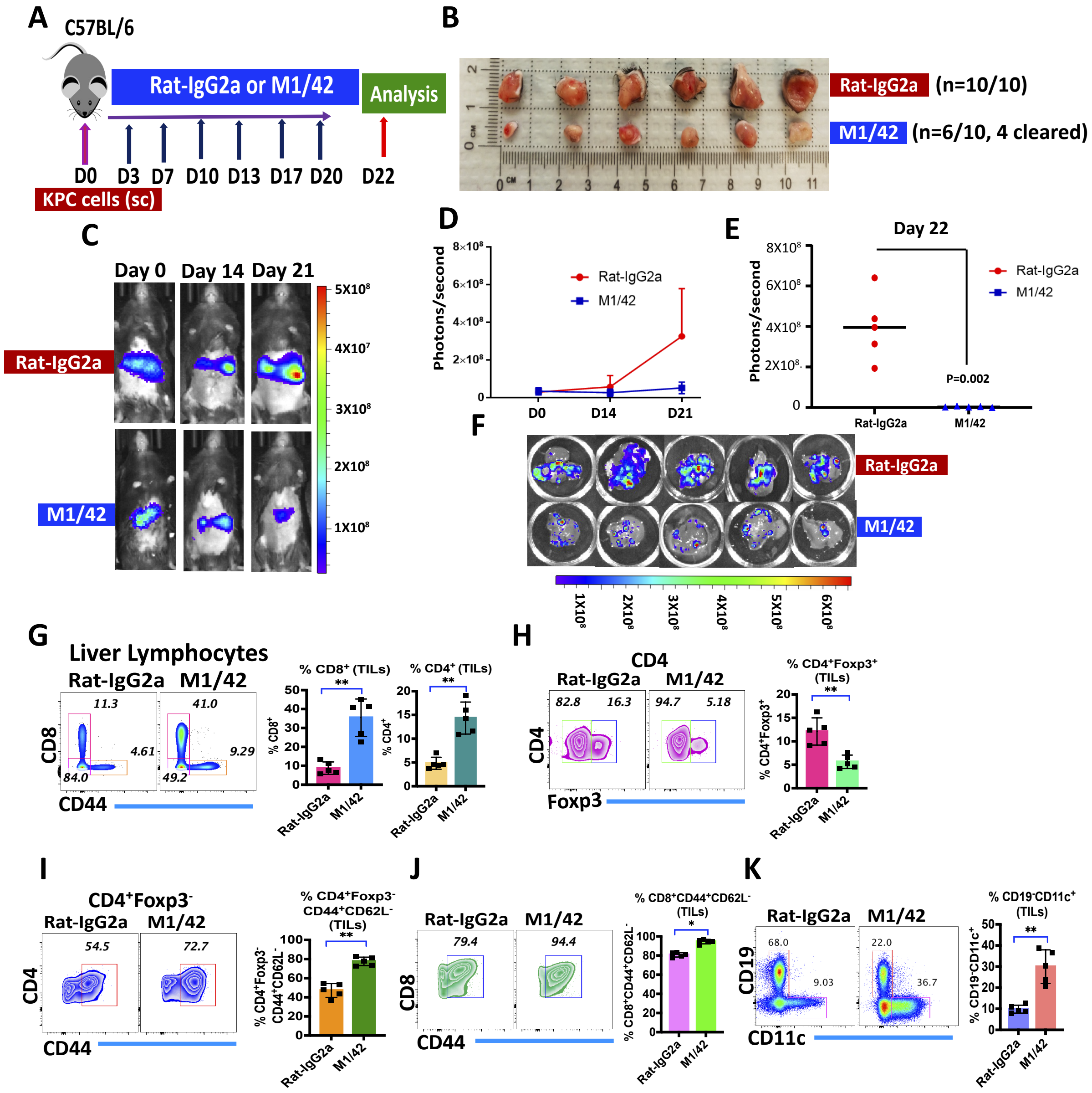
A
C57BL/6
Rat-IgG2a or M1/42
Analysis
D0 D3 D7 D10 D13 D17 D20 D22
KPC cells (sc)
B
Rat-IgG2a (n=10/10)
M1/42 (n=6/10, 4 cleared)
C
Day 0 Day 14 Day 21
Rat-IgG2a
M1/42
5X10^8
4X10^7
3X10^8
2X10^8
1X10^8
D
Photons/second
8×10^8
6×10^8
4×10^8
2×10^8
0
Rat-IgG2a
M1/42
D0 D14 D21
E
Day 22
Photons/second
8X10^8
6X10^8
4X10^8
2X10^8
0
Rat-IgG2a
M1/42
P=0.002
F
Rat-IgG2a
M1/42
1X10^8 2X10^8 3X10^8 4X10^8 5X10^8 6X10^8
G
Liver Lymphocytes
Rat-IgG2a M1/42
11.3 41.0
4.61 9.29
84.0 49.2
CD8
CD44
% CD8+ (TILs)
% CD8+
% CD4+ (TILs)
% CD4+
**
Rat-IgG2a M1/42
H
CD4
Rat-IgG2a M1/42
82.8 16.3 94.7 5.18
CD4
Foxp3
% CD4+Foxp3+ (TILs)
% CD4+Foxp3+
**
I
CD4+Foxp3-
Rat-IgG2a M1/42
54.5 72.7
CD4
CD44
% CD4+Foxp3- CD44+CD62L- (TILs)
% CD4+Foxp3- CD44+CD62L-
J
Rat-IgG2a M1/42
79.4 94.4
CD8
CD44
% CD8+CD44+CD62L- (TILs)
% CD8+CD44+CD62L-
*
K
Rat-IgG2a M1/42
68.0 9.03 22.0 36.7
CD19
CD11c
% CD19-CD11c+ (TILs)
% CD19-CD11c+
**

Fig. S3

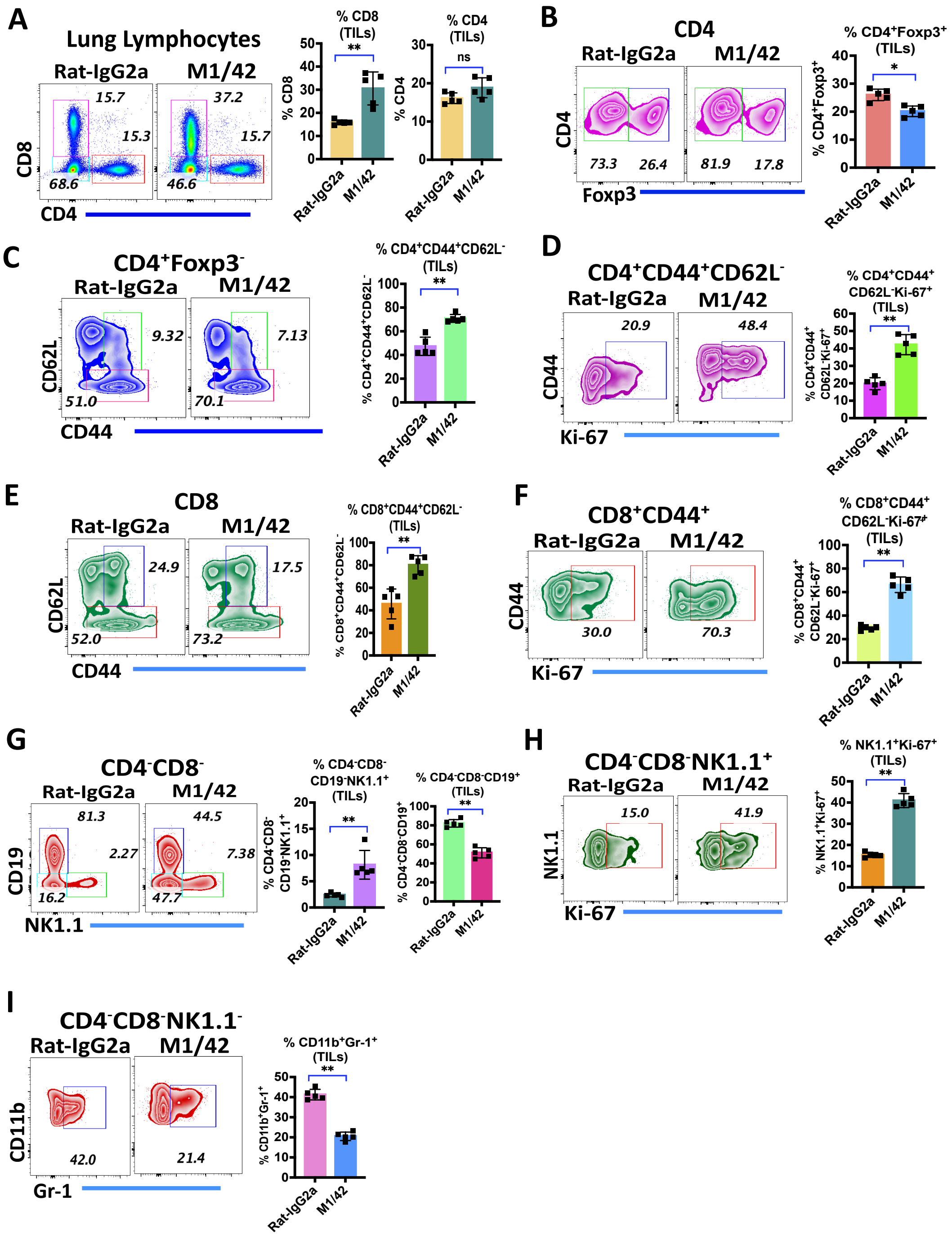

Fig. S4

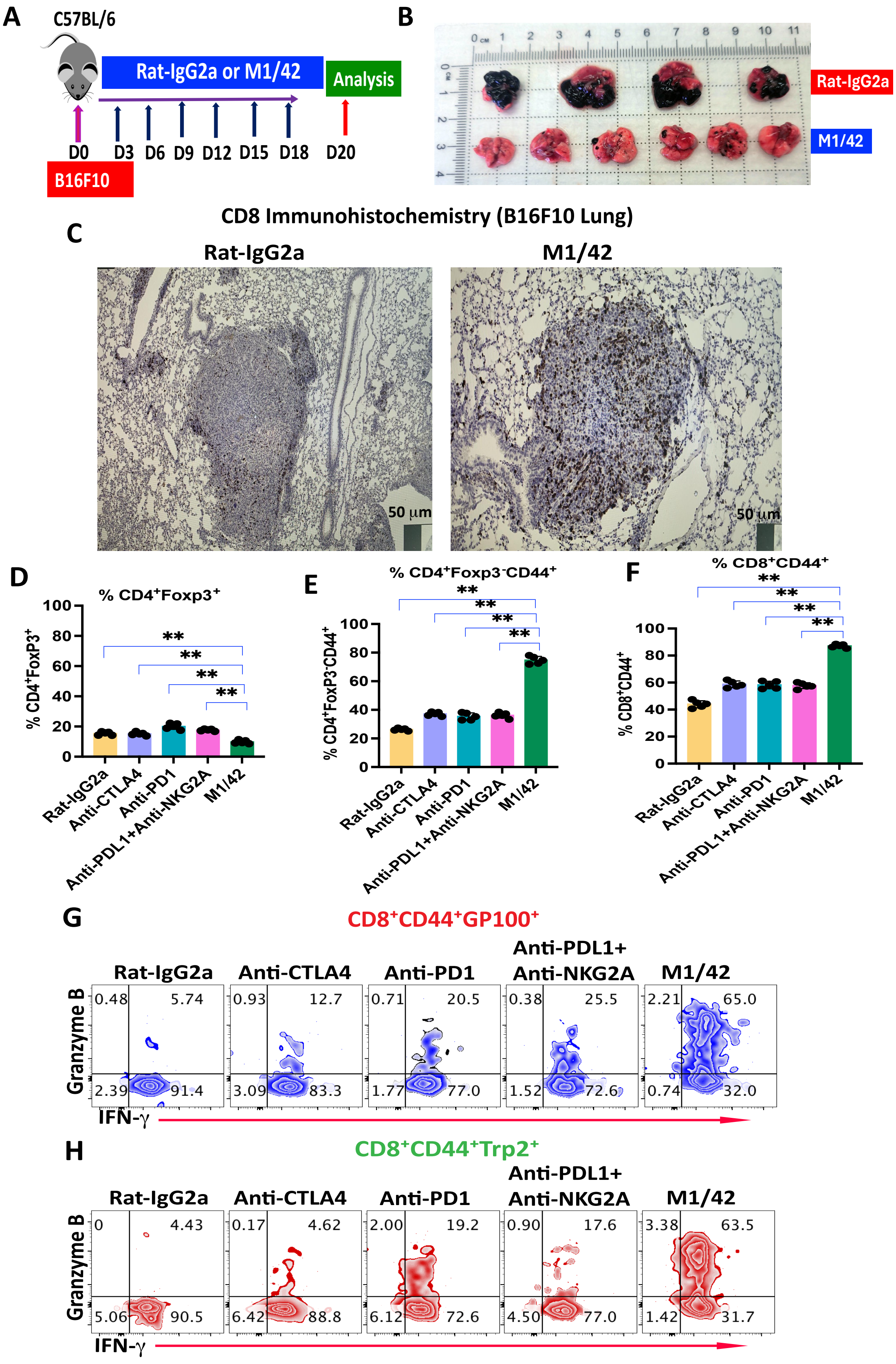
A
C57BL/6
Rat-IgG2a or M1/42
Analysis
D0 D3 D6 D9 D12 D15 D18 D20
B16F10
B
Rat-IgG2a
M1/42
C
CD8 Immunohistochemistry (B16F10 Lung)
Rat-IgG2a
M1/42
50 μm
D
% CD4+Foxp3+
% CD4+FoxP3+
E
% CD4+Foxp3-CD44+
% CD4+FoxP3-CD44+
F
% CD8+CD44+
% CD8+CD44+
Rat-IgG2a
Anti-CTLA4
Anti-PD1
Anti-PDL1+Anti-NKG2A
M1/42
G
CD8+CD44+GP100+
H
CD8+CD44+Trp2+
Granzyme B
IFN-γ

Fig. S5

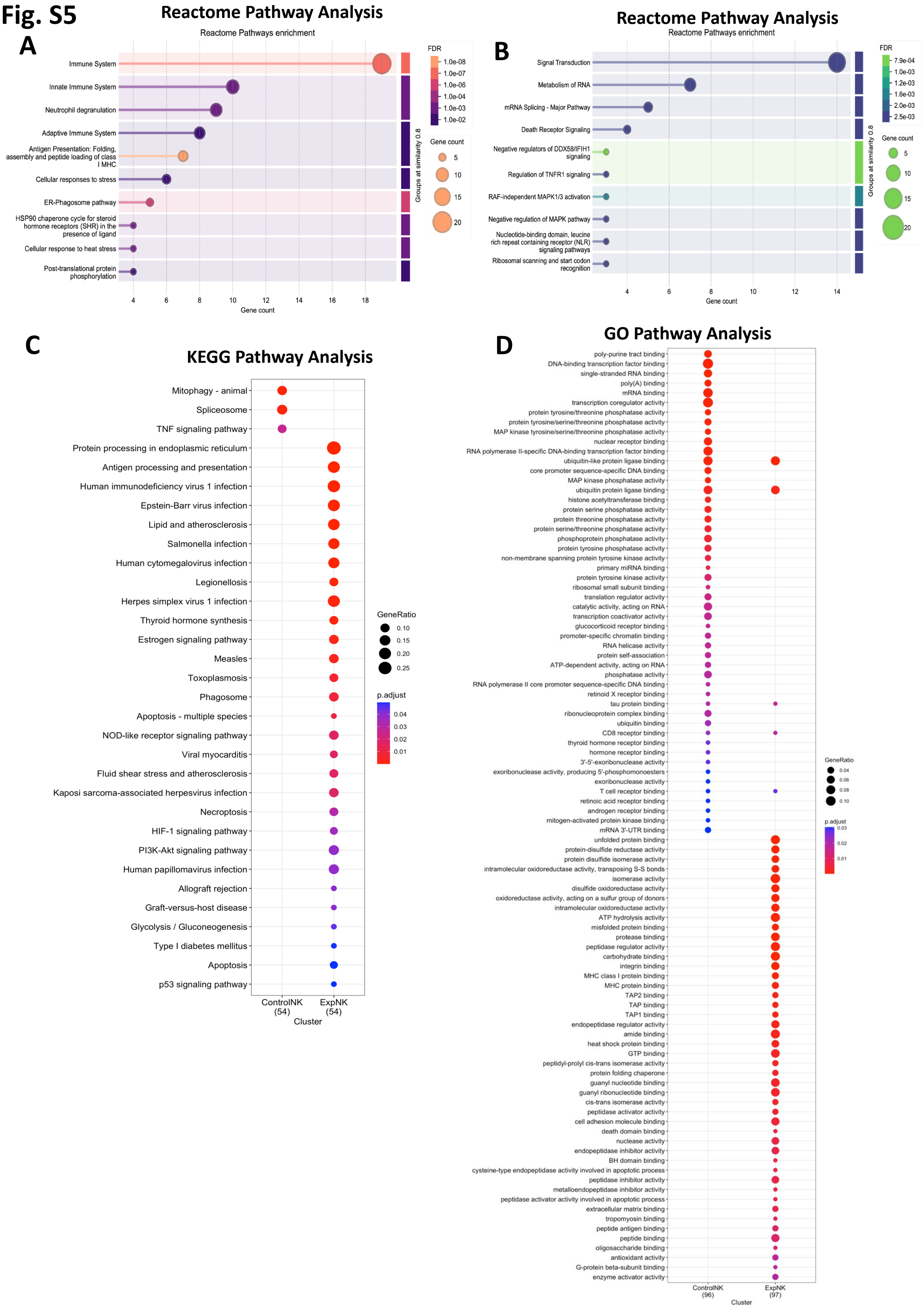
A
Reactome Pathway Analysis
Reactome Pathways enrichment
Immune System
Innate Immune System
Neutrophil degranulation
Adaptive Immune System
Antigen Presentation: Folding, assembly and peptide loading of class I MHC
Cellular responses to stress
ER-Phagosome pathway
HSP90 chaperone cycle for steroid hormone receptors (SHR) in the presence of ligand
Cellular response to heat stress
Post-translational protein phosphorylation
Gene count
Groups at similarity 0.8
FDR
1.0e-08
1.0e-07
1.0e-06
1.0e-05
1.0e-04
1.0e-03
1.0e-02
Gene count
5
10
15
20
B
Reactome Pathway Analysis
Reactome Pathways enrichment
Signal Transduction
Metabolism of RNA
mRNA Splicing - Major Pathway
Death Receptor Signaling
Negative regulators of DDX58/IFIH1 signaling
Regulation of TNFR1 signaling
RAF-independent MAPK1/3 activation
Negative regulation of MAPK pathway
Nucleotide-binding domain, leucine rich repeat containing receptor (NLR) signaling pathways
Ribosomal scanning and start codon recognition
Gene count
Groups at similarity 0.8
FDR
7.9e-04
1.0e-03
1.2e-03
1.6e-03
2.0e-03
2.5e-03
Gene count
5
10
15
20
C
KEGG Pathway Analysis
Mitophagy - animal
Spliceosome
TNF signaling pathway
Protein processing in endoplasmic reticulum
Antigen processing and presentation
Human immunodeficiency virus 1 infection
Epstein-Barr virus infection
Lipid and atherosclerosis
Salmonella infection
Human cytomegalovirus infection
Legionellosis
Herpes simplex virus 1 infection
Thyroid hormone synthesis
Estrogen signaling pathway
Measles
Toxoplasmosis
Phagosome
Apoptosis - multiple species
NOD-like receptor signaling pathway
Viral myocarditis
Fluid shear stress and atherosclerosis
Kaposi sarcoma-associated herpesvirus infection
Necroptosis
HIF-1 signaling pathway
PI3K-Akt signaling pathway
Human papillomavirus infection
Allograft rejection
Graft-versus-host disease
Glycolysis / Gluconeogenesis
Type I diabetes mellitus
Apoptosis
p53 signaling pathway
ControlNK (54)
ExpNK (54)
Cluster
GeneRatio
0.10
0.15
0.20
0.25
p.adjust
0.04
0.03
0.02
0.01
D
GO Pathway Analysis
poly-purine tract binding
DNA-binding transcription factor binding
single-stranded RNA binding
poly(A) binding
mRNA binding
transcription coregulator activity
protein tyrosine/threonine phosphatase activity
protein tyrosine/serine/threonine phosphatase activity
MAP kinase tyrosine/serine/threonine phosphatase activity
nuclear receptor binding
RNA polymerase II-specific DNA-binding transcription factor binding
ubiquitin-like protein ligase binding
core promoter sequence-specific DNA binding
MAP kinase phosphatase activity
ubiquitin protein ligase binding
histone acetyltransferase binding
protein serine phosphatase activity
protein threonine phosphatase activity
protein serine/threonine phosphatase activity
phosphoprotein phosphatase activity
protein tyrosine phosphatase activity
non-membrane spanning protein tyrosine kinase activity
primary miRNA binding
protein tyrosine kinase activity
ribosomal small subunit binding
translation regulator activity
catalytic activity, acting on RNA
transcription coactivator activity
glucocorticoid receptor binding
promoter-specific chromatin binding
RNA helicase activity
protein self-association
ATP-dependent activity, acting on RNA
phosphatase activity
RNA polymerase II core promoter sequence-specific DNA binding
retinoid X receptor binding
tau protein binding
ribonucleoprotein complex binding
ubiquitin binding
CD8 receptor binding
thyroid hormone receptor binding
hormone receptor binding
3'-5'-exoribonuclease activity
exoribonuclease activity, producing 5'-phosphomonoesters
exoribonuclease activity
T cell receptor binding
retinoic acid receptor binding
androgen receptor binding
mitogen-activated protein kinase binding
mRNA 3'-UTR binding
unfolded protein binding
protein-disulfide reductase activity
protein disulfide isomerase activity
intramolecular oxidoreductase activity, transposing S-S bonds
isomerase activity
disulfide oxidoreductase activity
oxidoreductase activity, acting on a sulfur group of donors
intramolecular oxidoreductase activity
ATP hydrolysis activity
misfolded protein binding
protease binding
peptidase regulator activity
carbohydrate binding
integrin binding
MHC class I protein binding
MHC protein binding
TAP2 binding
TAP binding
TAP1 binding
endopeptidase regulator activity
amide binding
heat shock protein binding
GTP binding
peptidyl-prolyl cis-trans isomerase activity
protein folding chaperone
guanyl nucleotide binding
guanyl ribonucleotide binding
cis-trans isomerase activity
peptidase activator activity
cell adhesion molecule binding
death domain binding
nuclease activity
endopeptidase inhibitor activity
BH domain binding
cysteine-type endopeptidase activity involved in apoptotic process
peptidase inhibitor activity
metalloendopeptidase inhibitor activity
peptidase activator activity involved in apoptotic process
extracellular matrix binding
tropomyosin binding
peptide antigen binding
peptide binding
oligosaccharide binding
antioxidant activity
G-protein beta-subunit binding
enzyme activator activity
ControlNK (96)
ExpNK (97)
Cluster
GeneRatio
0.04
0.06
0.08
0.10
p.adjust
0.03
0.02
0.01

### Fig. S6

Cluster 0 Comparison

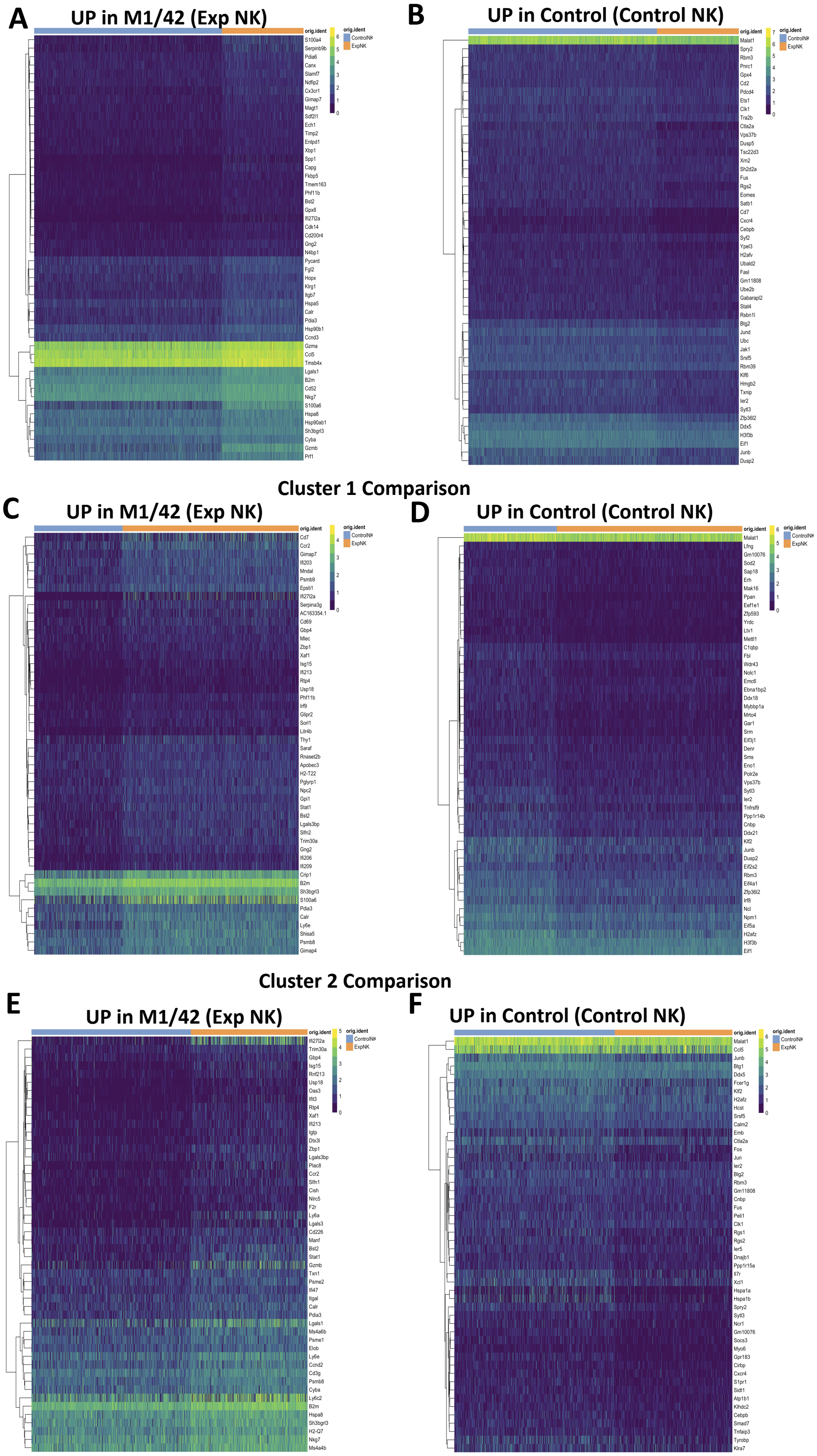

**Fig. S7**

### Cluster 0

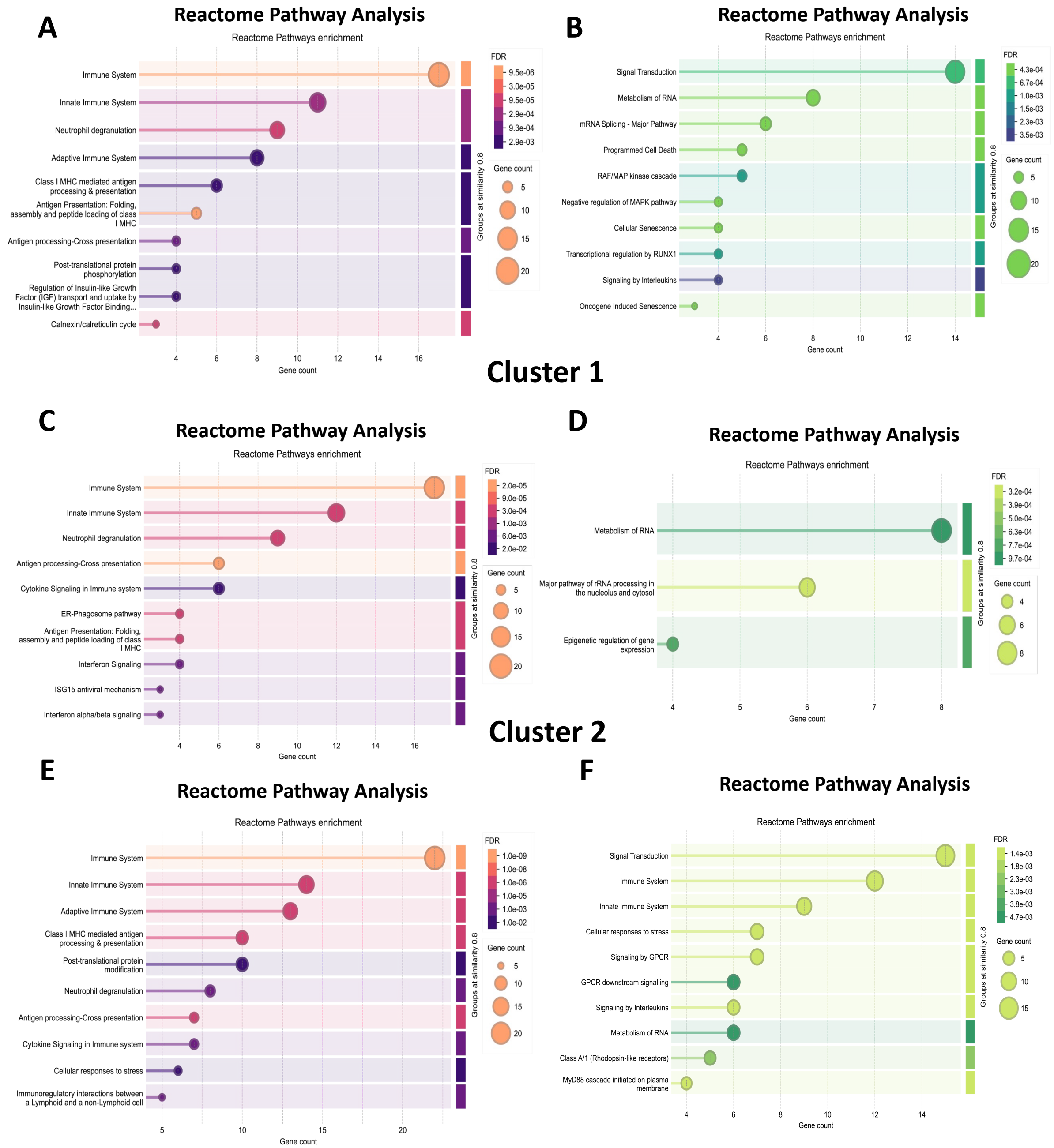

### Fig. S8

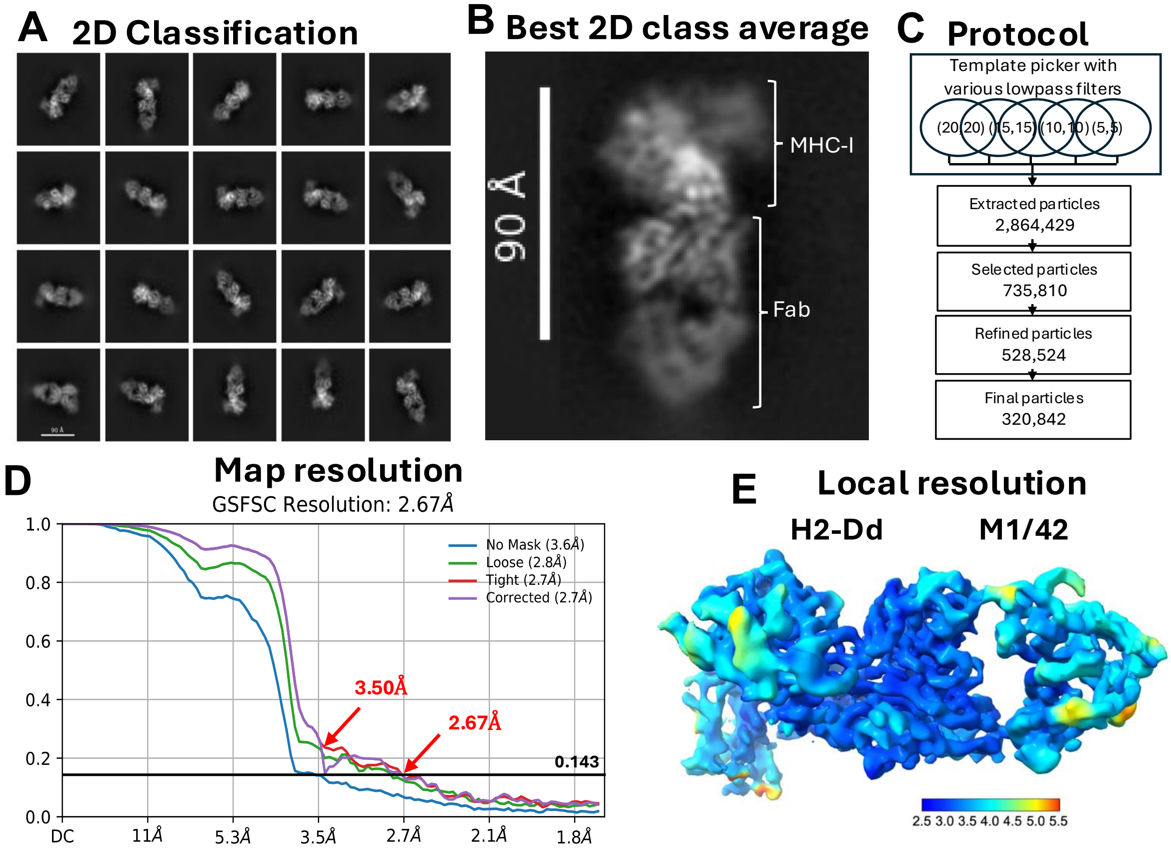


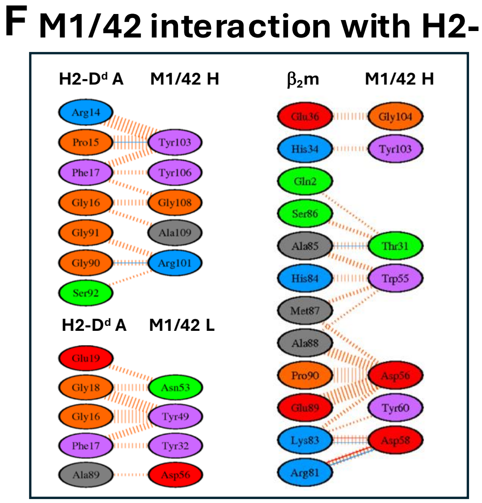


**G M1/42 contacts to H2-D$^d$**

| Res# | CH | Res# | CH | Dist. |
|---|---|---|---|---|
| ARG14 | A | TYR103 | H | 3.09 |
| PRO15 | A | TYR103 | H | 2.36 |
| GLY16 | A | GLY108 | H | 3.08 |
| GLY16 | A | ALA109 | H | 3.39 |
| PHE17 | A | TYR103 | H | 3.46 |
| PHE17 | A | GLY108 | H | 3.66 |
| PHE17 | A | TYR106 | H | 3.75 |
| GLY90 | A | ARG101 | H | 2.35 |
| GLY91 | A | ARG101 | H | 3.25 |
| SER92 | A | ARG101 | H | 3.77 |
| PHE17 | A | TYR32 | L | 3.50 |
| GLY16 | A | TYR49 | L | 3.46 |
| PHE17 | A | TYR49 | L | 3.29 |
| GLY18 | A | TYR49 | L | 3.28 |
| GLY16 | A | TYR49 | L | 3.60 |
| GLY18 | A | ASN53 | L | 3.18 |
| GLU19 | A | ASN53 | L | 3.20 |
| ALA89 | A | ASP56 | L | 3.70 |

| Res# | CH | Res# | CH | Dist. |
|---|---|---|---|---|
| GLN2 | B | THR31 | H | 3.68 |
| HIS34 | B | TYR103 | H | 3.50 |
| GLU36 | B | GLY104 | H | 3.20 |
| ARG81 | B | ASP58 | H | 2.67 |
| LYS83 | B | ASP56 | H | 3.24 |
| LYS83 | B | ASP58 | H | 2.35 |
| LYS83 | B | TYR60 | H | 3.83 |
| HIS84 | B | TRP55 | H | 2.99 |
| ALA85 | B | TRP55 | H | 3.62 |
| ALA85 | B | THR31 | H | 3.05 |
| SER86 | B | THR31 | H | 3.37 |
| MET87 | B | TRP55 | H | 3.08 |
| MET87 | B | ASP56 | H | 2.86 |
| ALA88 | B | TRP55 | H | 3.84 |
| ALA88 | B | ASP56 | H | 2.82 |
| GLU89 | B | ASP56 | H | 3.07 |
| PRO90 | B | ASP56 | H | 3.13 |

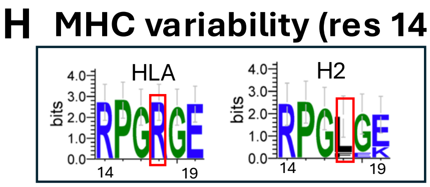


**I $\beta_2$m polymorphism at contacts**

| | | | | | | | |
|---|---|---|---|---|---|---|---|
| Human | D | N | V | T | L | S | Q |
| Mouse | H | K | A | S | M | A | Q |
| residue no | 34 | 83 | 85 | 86 | 87 | 88 | 89 |

**Table S1.** X-ray data collection and refinement statistics

| | M142+H2-D$^d$ |
|---|---|
| **PDBID** | **8TQ4** |
| **Data collection** | |
| Space group | $P4_322$ |
| Cell dimensions | |
| *a*, *b*, *c* (Å) | 121.55, 121.55, 327.20 |
| $\alpha$, $\beta$, $\gamma$ (°) | 90.0, 90.0, 90.0 |
| Resolution (Å)* | 95.57-3.59 (3.72-3.59) |
| $R_{sym}$ or $R_{merge}$ | 0.422 (1.857) |
| $I/\sigma(I)$ | 6.2 (1.3) |
| Completeness (%) | 95.0 (95.5) |
| Redundancy | 9.2 (9.4) |
| $R_{pim}$ | 0.139 (0.613) |
| $CC_{1/2}$ | 0.995 (0.803) |
| Twin | none |
| **Refinement** | |
| Resolution (Å)* | 95.57-3.59 (3.72-3.59) |
| No. unique reflections[§] | 28062 (2728) |
| $R_{work}/R_{free}$ (%) | 22.1/26.9 (27.8/31.8) |
| No. atoms | 12671 |
| Protein | 12671 |
| Water/ligands | 0/0 |
| B-factor Wilson/Average | 91.8/86.5 |
| Protein | 86.5 |
| Water/ligands | 0 |
| R.m.s. deviations | |
| bond length (Å) | 0.004 |
| bond angle (º) | 0.82 |
| Ramachandran | |
| favored (%) | 91.0 |
| allowed (%) | 7.0 |
| outliers (%) | 2.0 |
| Molprobity | |
| clashscore | 11.9 |

(*Values in parenthesis are for highest resolution shell, [§] Values in parenthesis are the number of reflections for $R_{free}$)

**Table S2.** Cryo-EM data collection, refinement and validation statistics

| | M1/42+H2-$D^d$ |
|---|---|
| EMDB ID | EMD-46600 |
| PDB-ID | 9D72 |
| **Data collection and processing** | |
| Magnification | 130,000 |
| Voltage (kV) | 300 |
| Electron exposure ($e^-$/Å) | 54.2 |
| Defocus range (μm) | -0.7 to -2.0 |
| Pixel size (Å/pixel) | 0.83 (binned) |
| Raw micrographs (no.) | 2,993 |
| Extract particles (no.) | 2,864,429 |
| Selected 2D particles (no.) | 735,810 |
| Refined particles (no.) | 528,545 (J244) |
| Particles for final map (no.) | 320,842 (J284) |
| Symmetry imposed | C1 |
| Map resolution (Å) | 2.67 (3.5 unmasked) |
| FSC threshold | 0.143 |
| **Refinement** | (J284) |
| Initial model | 8TQ4 |
| * Map sharpening B factor ($Å^2$) | 0 |
| Model composition | (rsr29.pdb) |
| Atoms | 6,040 |
| Residues | 790 |
| Ligands | 0 |
| Overall B-factor ($Å^2$) | |
| Protein (min/max/mean) | 36.8/314.5/146.2 |
| R.m.s. deviations | |
| bond length (Å) | 0.002 |
| bond angle (º) | 0.519 |
| CC (mask/volume/peaks) | 0.75/0.74/0.68 |
| Validation | |
| MolProbity score | 2.67 |
| Clash score | 13.4 |
| Ramachandran outliers (%) | 0.26 |
